# Thalamic nucleus reuniens preferentially targets inhibitory interneurons over pyramidal cells in hippocampal CA1 region

**DOI:** 10.1101/2021.09.30.462517

**Authors:** Lilya Andrianova, Paul J Banks, Clair A Booth, Erica S Brady, Gabriella Margetts-Smith, Shivali Kohli, Jonathan Cavanagh, Zafar I Bashir, Chris J McBain, Michael T Craig

## Abstract

The prefrontal – hippocampal – entorhinal system is perhaps the most widely-studied circuit in cognitive and systems neuroscience, due to its role in supporting cognitive functions such as working memory and decision-making. Disrupted communication within this circuit is a key feature of disorders such as schizophrenia and dementia. Nucleus reuniens (NRe) is a midline thalamic nucleus that sits at the nexus of this circuit, linking these regions together. As there are no direct projections from prefrontal cortex to hippocampus, the accepted model is that the NRe mediates prefrontal drive of hippocampal activity, although these connections are poorly defined at the cellular and synaptic level. Using *ex vivo* optogenetics and electrophysiology, alongside monosynaptic circuit-tracing, we sought to test the mechanisms through which NRe could drive hippocampal activity. Unexpectedly, we found no evidence that pyramidal cells in CA1 receive input from NRe, with midline thalamic input to hippocampus proper appearing selective for GABAergic interneurons. In other regions targeted by NRe, we found that pyramidal cells in prosubiculum and subiculum received synaptic inputs from NRe that were at least an order of magnitude weaker than those in prefrontal or entorhinal cortices. We conclude that, contrary to widely-held assumptions in the field, the hippocampal pyramidal cells are not a major target of nucleus reuniens.

## Introduction

Numerous aspects of cognition, such as memory [1] and decision-making [2], require the communication of information between prefrontal cortex (PFC), hippocampus (HPC), and entorhinal cortex (EC). Thalamic nucleus reuniens (NRe) sits at the nexus of this circuit and is implicated in conditions such a schizophrenia and epilepsy [3]. This circuit includes both direct connections between the cortical and hippocampal regions, as well as indirect routes via thalamus, making interpretation of the function of any individual component particularly challenging. Synchronous oscillations occur between PFC and HPC at theta frequency during decision-making [4,5], between EC and PFC at theta frequency during associative learning [6], and between EC and HPC at theta and gamma frequencies during spatial learning [7]. There is, however, an *anatomical anomaly* within this circuitry: despite the functional importance of harmonised activity, PFC does not project directly to HPC [see 8 for our recent confirmation]. The accepted view of this circuit is that NRe directly mediates communication between PFC and HPC [e.g. 9,10], which implicates NRe in cognitive functions such as working memory [11,12], and leading many to believe that the main function of NRe is to mediate prefrontal control over the hippocampus. However, NRe inactivation or lesion studies often fail to provide strong evidence for a role in spatial memory acquisition [13].

Anatomically, NRe forms reciprocal connections with PFC and subiculum, while uniquely for the thalamus, it also sends direct efferents to CA1; these efferents terminate alongside EC axons in stratum lacunosum-moleculare (SL-M) [14,15]. Hippocampal return projections to NRe arise entirely from subiculum, with neither dorsal nor ventral CA1 forming monosynaptic connections with neurons in NRe [16]. NRe sends projections to EC [17], but these are effectively not reciprocated [18] with perhaps only 2% of superficial EC neurons projecting to NRe [16]. Functionally, although electrical stimulation in NRe fails to elicit spiking in the CA1 pyramidal layer *in vivo* [19], it has been widely assumed that NRe targets CA1 pyramidal cells [11]. This assumption is reasonable as we are unaware of any cortical region in which thalamic inputs do *not* target pyramidal cells. Surprisingly, despite NRe’s important role in goal-directed spatial navigation [20] and working memory [21], NRe projections to CA1 have remained poorly defined. We previously reported that both NRe and entorhinal fibres terminate in stratum lacunosum-moleculare (SL-M) of CA1 where they target neurogliaform cells [22]. Optogenetic stimulation of NRe inputs to these neurogliaform cells elicits monosynaptic EPSCs that are defined by large NMDA receptor-mediated components.

Here, we sought to determine the nature of NRe inputs onto CA1 pyramidal cells. We hypothesised that a similarly large NMDA-R component in pyramidal cells to what we observed in neurogliaform cells could underlie observation that NRe-fEPSPs are greatly increased in CA1 during EC input coactivation [23]. However, we found no evidence direct monosynaptic projections from NRe to CA1 pyramidal cells, and only evidence of weak inputs to pyramidal cells in other areas of the hippocampal formation.

## Results

### NRe projections to hippocampal formation

We carried out stereotaxic injection of AAVs to express the light-sensitive channelrhodopsin2 variants Chronos or Chrimson [24] into NRe and prepared *ex vivo* brain slices in either coronal or horizontal planes to carry out patch clamp recordings targeting primarily prefrontal cortex and dorsal hippocampus, or entorhinal cortex and ventral hippocampus, respectively (example distribution of NRe axons in hippocampus shown in Fig. S1). We subdivided the hippocampus into CA1, prosubiculum and subiculum, using the Lorente de Nó boundaries [25]. We denoted the boundary of CA1 to be where the dense, ordered nature of stratum pyramidale ended, with prosubiculum continuing in an area with a slight laminar-like organisation in the superficial region (see figure 2 and particularly figure 11 of the Lorente De Nó paper), with subiculum beginning when no obvious laminar structure was apparent through DIC optics. The boundaries that we used correspond well with those reported by more recently [26], and are shown in Fig. 1A (ventral hippocampus) and Fig. 2A (dorsal hippocampus).

**Figure 1:**
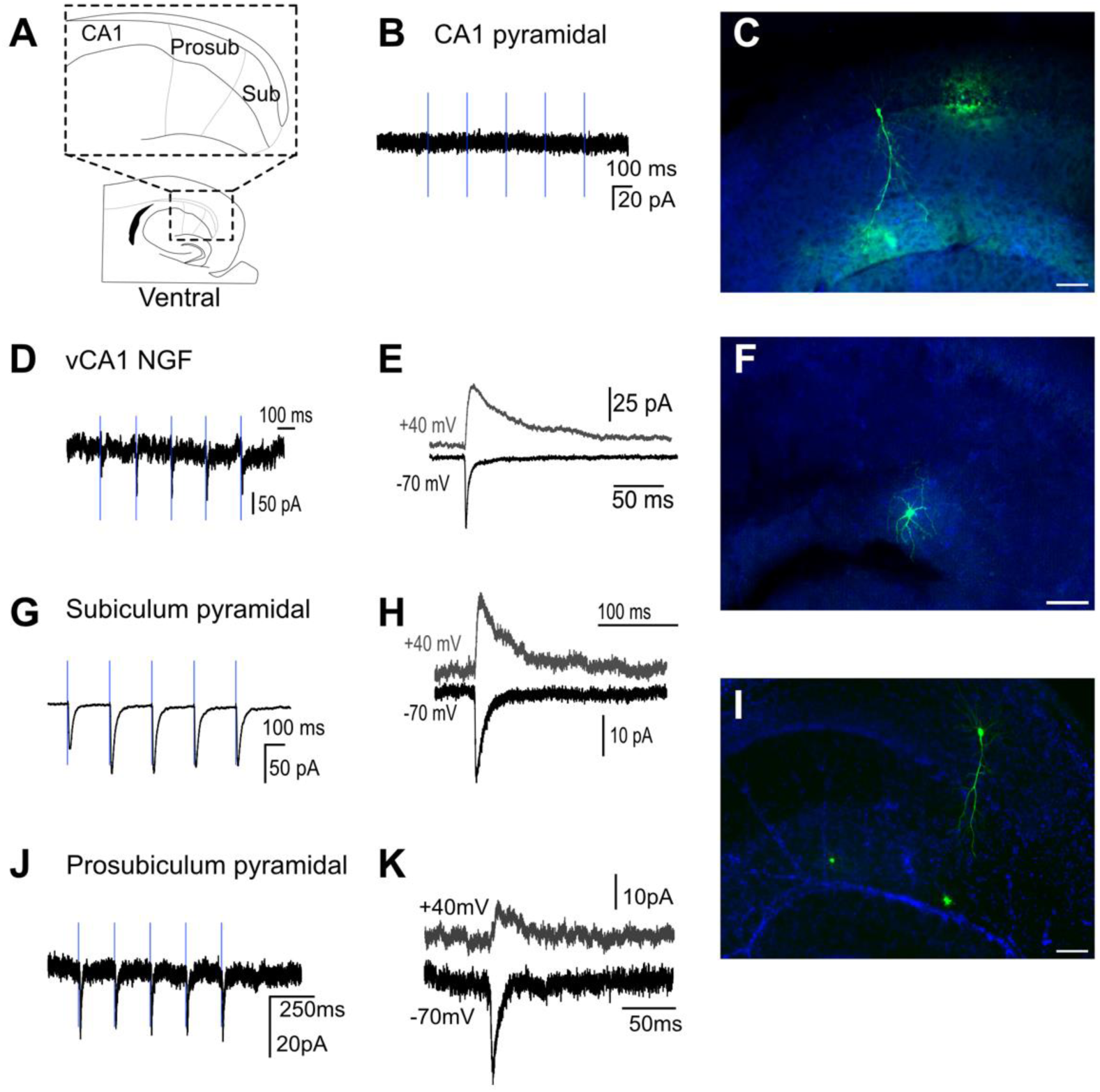
Prosubiculum pyramidal cells, unlike those in ventral CA1, do receive an input from NRe. Schematic representation of boundaries between dorsal CA1, prosubiculum and subiculum in horizontal orientation. **A**, CA1 / prosubiculum boundary denoted as region where additional pyramidal cell layers arise ‘below’ tightly-packed layer that is a continuation of CA1 stratum pyramidale, after Lorento de No (1934). **B,** no response recorded from dCA1 pyramidal cell. **C,** *post hoc* recovery of a non-responsive pyramidal cell in dCA1. **D,** representative response trace from a ventral NGF. **E,** maximal AMPA and NMDA components of the responsive ventral NGF cell. **F,** A post hoc recovery from a ventral NGF cell. **G,** response trace from responsive pyramidal cell in dorsal subiculum. **H,** maximal AMPA and NMDA components of a responsive pyramidal cell from subiculum. **I,** post hoc recovery of a responsive pyramidal neuron in prosubiculum. **J,** a response trace from a responsive pyramidal cell in dorsal prosubiculum. **K,** maximal AMPA and NMDA components of a responsive pyramidal cell from prosubiculum. Scale bar 100 um.

**Figure 2:**
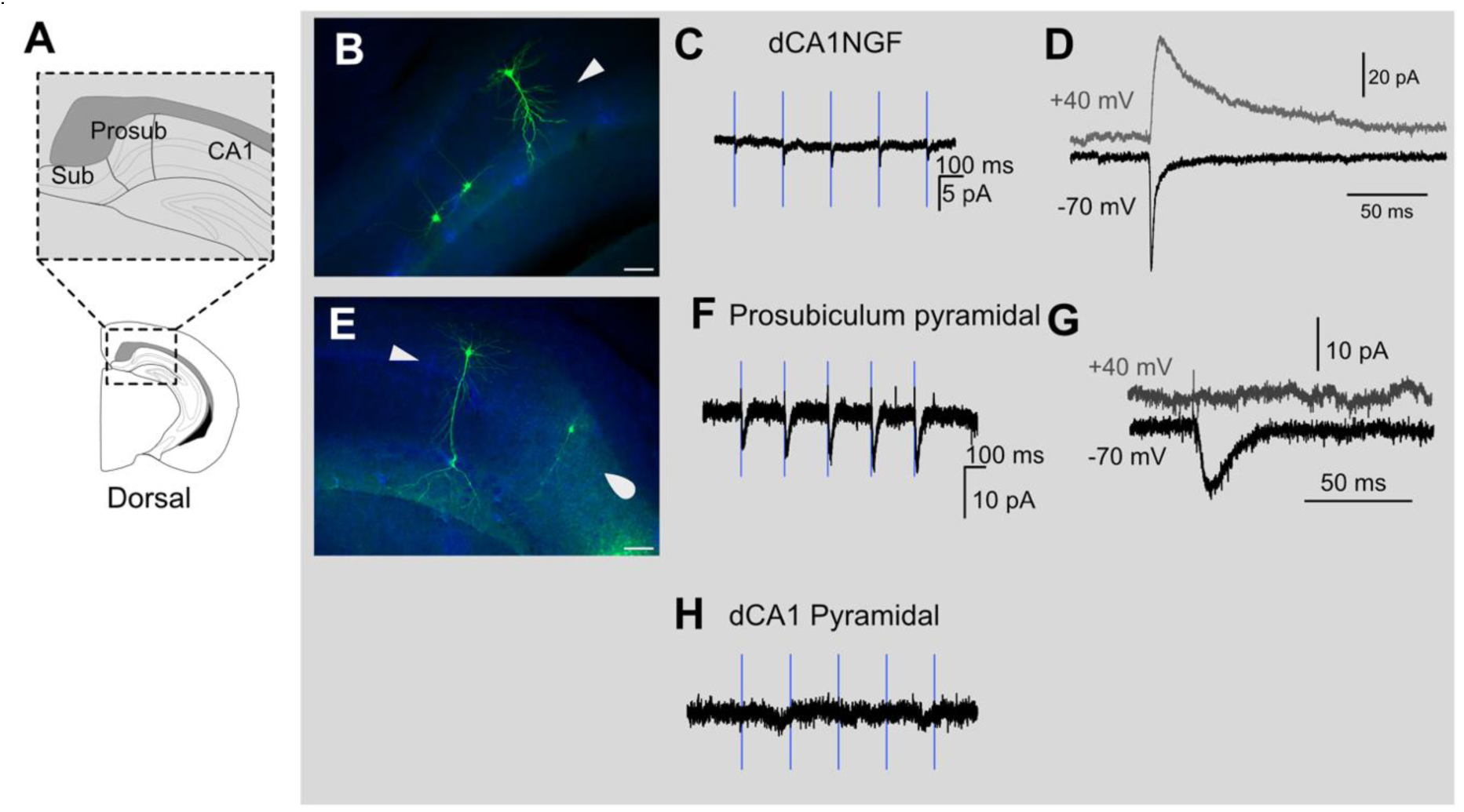
Prosubiculum pyramidal cells, unlike those in dorsal CA1, do receive an input from NRe. **A,** Schematic representation of boundaries between dorsal CA1, prosubiculum and subiculum in coronal orientation. **B,** A *post hoc* recoveries of two neurogliaform cells and a non-responding CA1 pyramidal cell. **C**, A representative response trace from a dorsal NGF. **D,** Maximal AMPA and NMDA components of a responsive NGF from dorsal CA1**. E,** *Post hoc* recoveries of two pyramidal cells, the non-responding CA1 pyramidal cell (labelled by arrowhead) and responding pyramidal cell in the prosubiculum (labelled with rounded arrowhead) as well as a dorsal NGF. **F,** A response trace from a responsive pyramidal neuron from dorsal prosubiculum. **G,** Maximal AMPA and NMDA components of a responsive pyramidal neuron from dorsal prosubiculum. **H,** A response trace from a non-responsive pyramidal neuron from dorsal CA1. Scale bar 100 um.

As NRe sends a substantially denser projection to ventral over dorsal hippocampus [14], we first carried out patch clamp recordings in ventral CA1 (vCA1). We found that optogenetic activation of NRe axons in CA1 failed to elicit post-synaptic EPSCs in vCA1 pyramidal cells (Fig. 1B & 3A). We observed the same result in dorsal CA1 (dCA1) (Fig. 2B & 3A). In each slice tested, we only recorded a pyramidal cell as non-responsive when we could evoke a post-synaptic response in neurogliaform cells that, with their high NRe input probability acted as positive controls (Fig. 1D – F for vCA1 and Fig. 2C – E for dCA1; summary data Fig. 3A). We found that there was no difference in the magnitude of NRe-EPSC on neurogliaform cells when evoked using either Chronos or Chrimson (Fig. S2). Additionally, we found no differences in NRe input probability or synaptic strength between dorsal and ventral hippocampus, nor did we observe differences in NRe-EPSC magnitude between MGE- and CGE-derived neurogliaform cells (Fig. S3).

**Figure 3:**
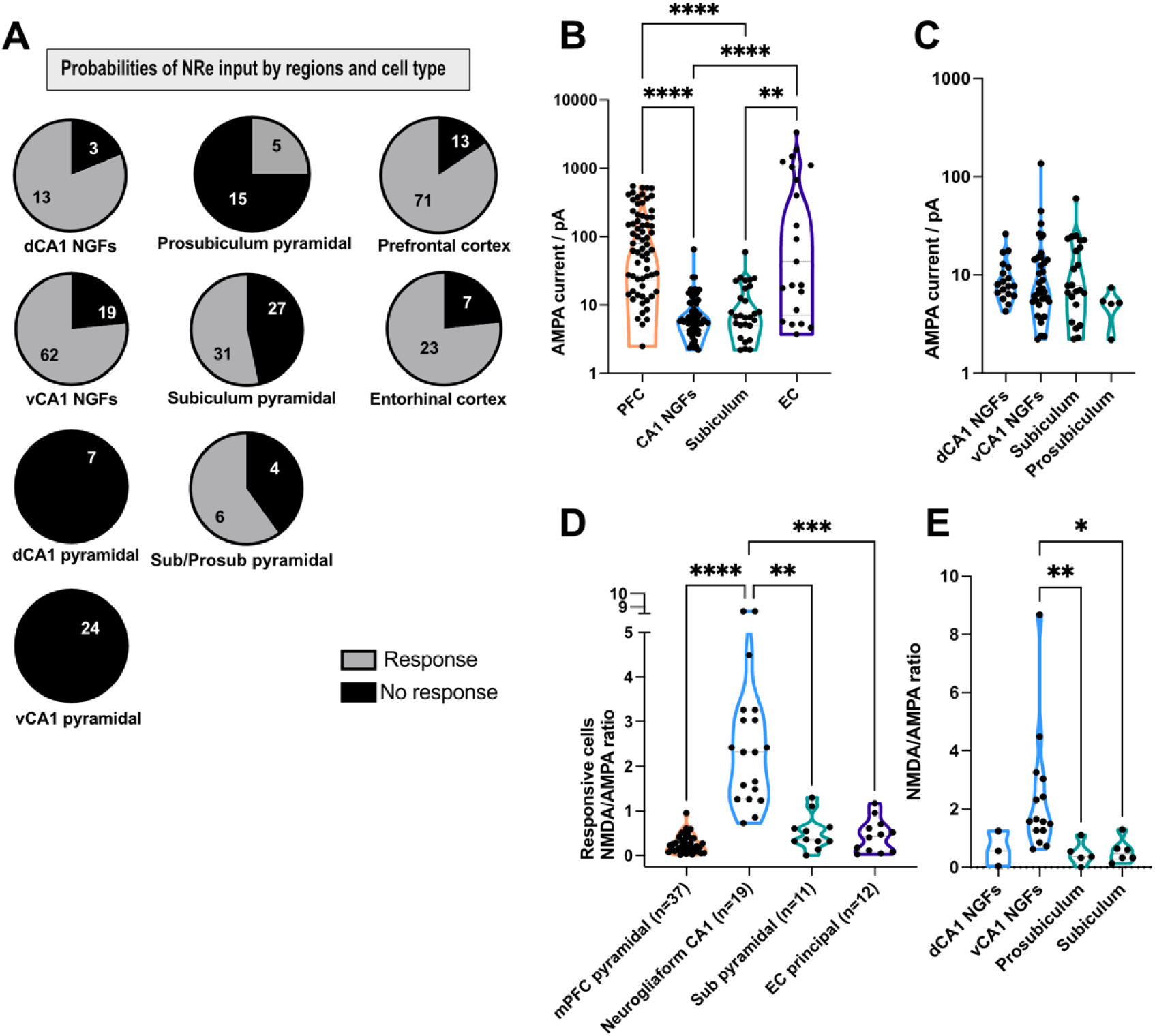
Summary data for all different cell types in the hippocampal formation and cortical areas. **A,** NRe input probability for cells patched in hippocampus: dorsal NGFs (81.3%) and ventral NGFs (76.5%), dorsal (0%) and ventral (0%) CA1 pyramidal cells, prosubiculum (66.7%), subiculum (46.6%) and undetermined prosubiculum/subiculum (60%) pyramidal neurons, prefrontal cortex (84.5%) and entorhinal cortex (76.7%). **B,** AMPA currents vary across different NRe target regions, with average AMPA currents being −142±19 pA for PFC pyramidal cells, −9.1±1.2 pA for NGFs, −12±2.3 pA for pyramidal cells in subiculum and -551±190 pA for principal cells in the entorhinal cortex. Kruskal-Wallis test (with Dunn’s multiple comparison test) showed significant differences between CA1 NGFs vs EC (p<0.0001), CA1 NGFs vs PFC (p<0.0001), EC vs Subiculum (p=0.0014) and Subiculum vs PFC (p<0.0001). **C,** AMPA currents across different regions with the average being −9.88±1.3 pA (dCA1 NGFs), −14.33±3.7pA (vCA1 NGFs), −13.32±2.7 pA (Subiculum pyramidal cells), -5.06±0.8 pA (Prosubiculum pyramidal cells), no significant difference found using a one-way ANOVA with Tukey’s multiple comparisons test. **D,** NMDA/AMPA ratios of responding cells in different regions, average ratios were 1.1±0.26 (dCA1 NGFs), 3.8±1.7(vCA1 NGFs), 0.47±0.18 (Subiculum pyramidal cells), 0.55±0.17 (Prosubiculum pyramidal cells). Using a Kruskal-Wallis test with Dunn’s multiple comparisons test significant different were found for NGFs vs Subiculum pyramidal cells (p=0.0047), EC principal vs NGFs (p=0.0003) and PFC pyramidal cells vs NGFs (p<0.0001). **E,** Kruskal-Wallis test with Dunn’s test for multiple comparisons, vNGFs cells have a significantly higher NMDA/AMPA ration when compared to subiculum and prosubiculum pyramidal neurons (p=0.0206 and p=0.0074, respectively). N = cell.

Unlike CA1, we found that NRe projections to other parts of the hippocampus (prosubiculum and subiculum; Fig. 1G – K for vCA1 and Fig. 2F – H for dCA1) did indeed target principal cells although the synaptic strength of these inputs was low (-13.32±2.7 pA for subiculum; summary data in Fig. 3), suggesting that NRe projections to hippocampal principal cells are either absent (CA1) or subthreshold (prosubiculum and subiculum Fig. 3), and unlikely to elicit firing. NRe projections to CA1 neurogliaform cells had a comparable magnitude of the AMPA- receptor mediated component of the EPSC (Fig. 3C) but the higher input resistance of inhibitory interneurons combined with the larger NMDA-AMPA ratio (Fig. 3E) would suggest that NRe could drive neurogliaform cells to spike, perhaps during periods of ongoing network activity.

### NRe projections to non-hippocampal areas

We also investigated the connectivity of the NRe to the cortical target regions, prefrontal (Fig. 4) and entorhinal cortices (Fig. 5). Similarly to the hippocampal region, the non-responding cells were only taken into the account if the same slice contained a positive control. Directly comparing NRe-evoked EPSC responses in cortical and hippocampal neurons, we found that principal cells in MEC and PFC pyramidal cells displayed significantly larger AMPA receptor-mediated EPSCs than subiculum pyramidal cells or CA1 NGFs (Fig. 3A, B & D). These data would suggest that, from a functional perspective, EC and PFC, and not the hippocampus, may be the principal targets of thalamic nucleus reuniens.

**Figure 4:**
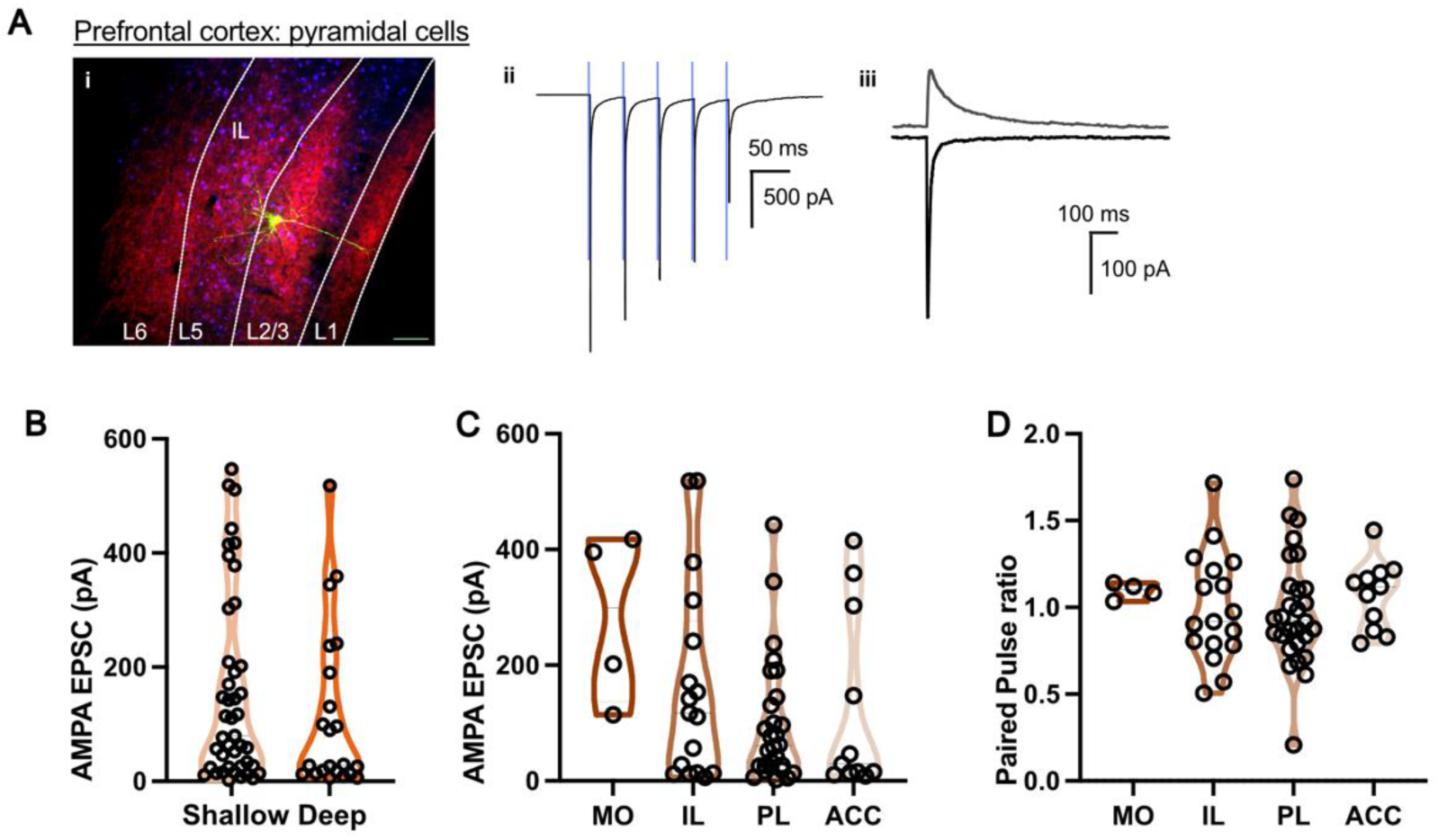
NRe-EPSCs are consistent across all subdivisions of the prefrontal cortex. **A,** A representative *post hoc* recovery of a pyramidal neuron in shallow layers of prefrontal cortex (**i**, scale bar 100-micron), optogenetic response trace (**ii**), AMPA and NMDA currents (**iii**). **B**, AMPA-R mediated NRe-EPSC did not vary significant between deep (layer 2 to 3) and shallow (layer 5 to 6) pyramidal cells in PFC (shallow *vs* deep: 155 ± 25 pA (n=43) *vs* 114 ± 30 pA (n=22); p=0.232, Mann-Whitney test). **C**, AMPA-R mediated NRe-EPSC did not vary between different subdivisions of PFC (MO *vs* IL *vs* PL *vs* ACC: 282 ± 74 pA (n=4) *vs* 165 ± 42 pA (n=17) *vs* 98 ± 20 pA (n=28) *vs* 147 ± 56 pA (n=9 cells); p=0.46, one-way ANOVA). **D,** paired pulse ratio of AMPA-R mediated NRe-EPSC did not vary between different subdivisions of PFC (MO *vs* IL *vs* PL *vs* ACC: 1.1 ± 0.02 *vs* 1.0 ± 0.08 *vs* 0.98 ± 0.06 *vs* 1.1 ± 0.06; p=0.15, one-way ANOVA). N = cells.

**Figure 5:**
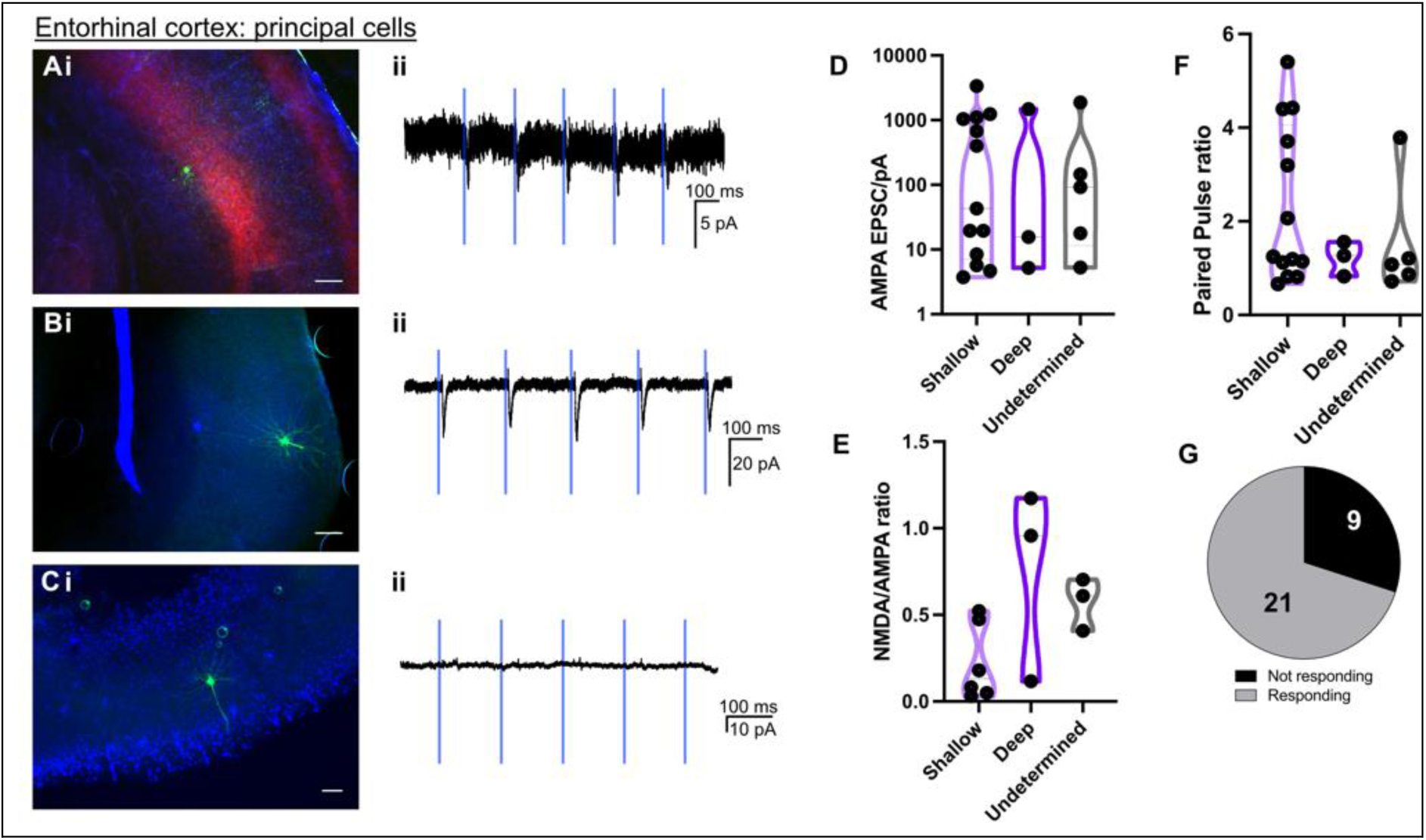
NRe-EPSCs are consistent across shallow and deep layers of entorhinal cortex. **A***, Post hoc* recovery of layer 5 mEC pyramidal neuron (**i**) and optogenetic response trace (**ii**). **B**, *Post hoc* recovery of layer 2 mEC stellate cell (**i**) and optogenetic response trace (**ii**). **C**, *Post hoc* recovery of layer 2 mEC pyramidal neuron (**i**) and lack of response to optogenetic stimulus (**ii**). **D**, AMPA-R mediated NRe-EPSCs are not different in shallow vs deep layers of mEC. **E**, NMDA/AMPA ratio in responsive neurons in shallow and deep layers of mEC is not significantly different. **F**, Paired pulse ratio in responsive neurons in shallow and deep layers of mEC is not significantly different. **G**, Input probability of mEC neurons.

When looking at the prefrontal cortex overall, the majority of the pyramidal cells were responsive to the optogenetic stimulation (Fig. 3A & 4A). We saw a similar result in medial EC (Fig. 3A & 5A). We saw no difference in NRe-EPSC amplitude in PFC when comparing NRe input between medial orbital, infralimbic, prelimbic, and anterior cingulate areas (Fig. 4B-D). While there are at least two main principal cell subtypes in EC [27–29], we did not parse EC neurons further into stellate or pyramidal cells so cannot comment on whether the large variability in NR-EPSC magnitude observed (Fig. 5D) corresponded to different subtypes of neuron. Nevertheless, these data confirm that NRe forms strong synaptic inputs onto principal neurons in both prefrontal and entorhinal areas.

To exclude the possibility that a lack of electrophysiological response from NRe axons onto CA1 pyramidal cells was unique to mouse, we undertook an independent replication of this finding in a different laboratory using rats. We found that 0/9 dorsal CA1 pyramidal cells and 0/17 ventral CA1 pyramidal cells in rat hippocampus displayed post-synaptic responses in response to optogenetic stimulation fo NRe axons, but that more than half of all interneurons in SL-M did indeed receive a response (Figure S4). Importantly, prefrontal cortex pyramidal neurons from the same preparations responded robustly to NRe input with similar currents to those seen in mouse [30].

### Monosynaptic retrograde tracing from CA1 pyramidal cells

Although our patch-clamp electrophysiological recordings failed to find evidence of somatic EPSCs mediated by NRe inputs in CA1 pyramidal cells, given that NRe ESPCs in neurogliaform cells and subiculum pyramidal cells have a small magnitude (Fig. 3) we could not exclude the possibility that inputs were present but undetectable due to dendritic filtering of the NRe inputs arriving in the distal region of apical dendrites, despite having found no evidence of silent NMDA-R only synapses. Consequently, we carried out rabies-assisted monosynaptic circuit tracing in *Emx1-cre* mice that conditionally expressed the avian TVA receptor only in pyramidal cells (see methods & Fig. 6). In our experimental approach, as TVA was expressed genetically in pyramidal cells, it was possible for rabies viruses to locally transduce pyramidal cells in CA1. However, we used a pseudotyped glycoprotein-deficit SAD B19 version of the rabies virus that is unable to replicate and spread from transduced neurons to express mCherry. Prior to rabies injection, we used an AAV helper virus to express GFP and the rabies glycoprotein (see methods and Figure S5). Thus, in our experiment, double-labelled GFP and mCherry cells in CA1 or subiculum are starter cells from which the rabies virus could retrogradely spread a single synapse, while mCherry-only cells in CA1 could be either presynaptic neurons or transduced with the pseudotyped vector. mCherry-only cells in other brain regions would be those presynaptic to CA1 pyramidal cells. Some data from this tracing experiment has previously been published elsewhere [8].

**Figure 6:**
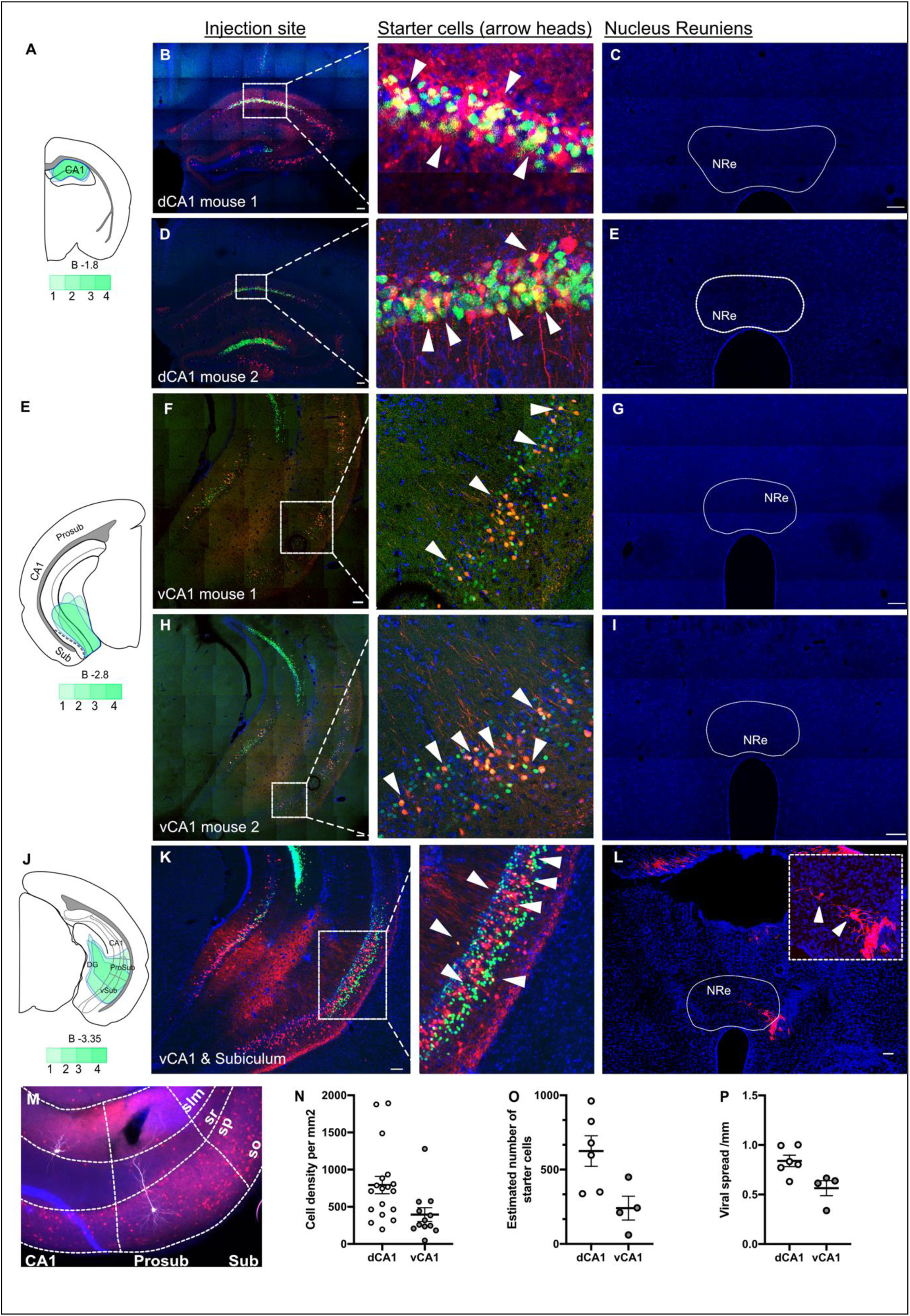
Monosynaptic retrograde tracing from hippocampus proper shows no cells originating from NRe, unless the starter cells are present in subiculum region. **A,** Injection site for dorsal CA1 and viral spreads for the helper virus (green cells). **B&D**, Injection site of a dCA1 injection: green cells contain helper virus and red cells are pseudotyped rabies virus positive and starter cells (yellow) shown closer on the inset, shown by arrowheads. **C & E,** No cells found in any thalamic section containing NRe. **E**, Injection site for ventral CA1 and viral spreads for the helper virus (green cells). **F & H,** Injection site of a vCA1 injection: green cells contain helper virus and red cells are pseudotyped rabies virus positive and starter cells (yellow) shown closer on the inset, shown by arrowheads. **G & I,** No cells found in any thalamic section containing NRe. **J**, Injection site for ventral CA1&subiculum and viral spreads for the helper virus (green cells). **K,** Injection site of a vCA1 injection with virus spreading to subiculum: green cells contain helper virus and red cells are pseudotyped rabies virus positive and starter cells (yellow) shown closer on the inset, shown by arrowheads. **L**, Cells and fibres seen in NRe as well as the fornix. **M**, A post hoc recovery of pyramidal cell in prosubiculum and neurogliaform cell in SLM layer of CA1 in horizontal slice orientation; boundary between CA1 and prosubiculum can be determined by reduction in SR layer thickness and reduction in canonical cell density in SP layer and the boundary between prosubiculum and subiculum is where SR layer almost disappears. **N**, Cell density showing number of cells containing the pseudo-typed rabies virus in dorsal and ventral CA1 regions. **O,** Estimated number of starter cells, calculated from cell density and viral spread. **P,** Viral spread distance in dorsal and ventral CA1 regions. For more details on retrograde monosynaptic tracing experiment data see [8].

We failed to find evidence of monosynaptic NRe inputs to pyramidal cells in either dorsal (Fig. 6A & E; n=6 mice) or ventral (Fig. 6 F & I, n=4) CA1 but, as expected, we saw consistent retrograde labelling in entorhinal cortices, medial and lateral septa and subiculum (we previously published some of this dataset in [8], confirming prior findings from Sun and colleagues [31]). We only observed retrograde labelling of NRe neurons when injections into ventral hippocampal formation included starter cells in both vCA1 and ventral prosubiculum / subiculum (Fig. 6J-L; n=3; all mice had retrogradely-labelled cells in NRe). The boundaries between the CA1, prosubiculum and subiculum regions in ventral hippocampus in horizontal plane are shown in Fig. 6M. Quantification of double-labelled starter cells, estimated overall number of starter cells as well as viral spread data is shown in Fig.6 N-P, and reproduced from our previous work [8]. To exclude the possibility that *Emx1* is not ubiquitously expressed throughout the hippocampus, we carried out a secondary analysis of RNAseq datasets publicly available from [32] and found that *Emx1* is indeed expressed throughout CA1 (Fig. S6).

## Discussion

Anatomical studies of NRe connectivity have led many to believe that the main function of NRe is to mediate prefrontal control over the hippocampus [e.g. 11]. However, our findings challenge this core assumption. We found that the magnitude of synaptic currents driven by NRe axons is at least an order of magnitude greater in EC and PFC than in either CA1 or subiculum. Furthermore, we have found that pyramidal cells in CA1 appear *not* receive monosynaptic inputs from NRe (determined using both anterograde optogenetic circuit-mapping and monosynaptic retrograde rabies tracing). It appears that NRe preferentially targets inhibitory interneurons in CA1 and pyramidal cells in prosubiculum.

### Do CA1 pyramidal cells receive input from NRe?

Our data cannot entirely exclude the possibility of a monosynaptic projection from NRe to CA1 pyramidal cells, but if the connection does exist then it must be very sparse. Unlike hippocampus proper, we found that pyramidal cells in both prosubiculum and subiculum do indeed receive an input from NRe. The lack of input to pyramidal cells in our study was very specific to CA1; pyramidal cells in prosubiculum, identified using Lorente de Nó anatomical definitions [25], did receive direct input from NRe. While some regard prosubiculum as the distal region of CA1, recent transcriptomic studies confirm that it is genetically distinct from both CA1 and subiculum [33], and our findings support this. The regions that we recorded from in prosubiculum were clearly outwith the boundaries of CA1, where the dense organisation of neurons in stratum pyramidale; no pyramidal neuron recorded within the more highly-ordered CA1 responded to optogenetic stimulation of thalamic axons, while we always used neurogliaform cells as positive controls. It has been shown that NRe selectively innervates pyramidal cells in ventral subiculum that target nucleus accumbens and lateral habenula, whilst avoiding those that target PFC [34].

When we first reported these findings as a preprint in late 2021 [35], the results appeared controversial. Indeed, while a recent publication reported a direct monosynaptic projection from NRe to dorsal CA1 pyramidal cells [36], the key recordings in that study appear to have been in the subiculum and/or prosubiculum areas of the hippocampus rather than adjacent CA1 (see Figure 1 of this study - the only figure showing recording location). However, other published studies would provide support for our findings. An ultrastructural study reported that synapses from thalamic axons in CA1 make significantly fewer contacts per dendrite than those from entorhinal axons [37], supporting our electrophysiological data and the hypothesis that NRe input to hippocampus serves to modulate, rather than drive, HPC activity, perhaps via inhibitory tone in SL-M. This ultrastructural study found projections from NRe onto spiny dendrites and thus concluded that these contacts were onto CA1 pyramidal cells as GABAergic interneurons are typically considered aspiny. However, recent work has demonstrated the existence of a population of spiny parvalbumin-expressing interneurons in CA1 that have dendrites in SL-M [38], which may have been the target of these NRe neurons. Another recent study reported close proximity of NRe axons onto PSD95-expressing dendritic spines in CA1, again concluding that these projections were onto pyramidal cells [39]. Again, while PSD95 is typically considered a marker of glutamatergic neurons, PSD95 has long been known to also exist on GABAergic neurons, particularly Erb4-expressing parvalbumin cells, as found by ourselves and others [e.g. 40–42].

A recent article from Leprince and colleagues [43] confirmed our main finding using a combination of *in vivo* and *ex vivo* approaches: they found that EC inputs strongly drive CA1 network activity in early postnatal circuitry and that, while NRe stimulation could drive network activity in CA1, this is almost entirely through inhibitory mechanisms. Indeed, this study reported that only 2 out of 26 CA1 pyramidal cells received input from NRe and it is unclear whether this connection persists into adulthood, with the authors speculating that it may be a transitory circuit in development that helps ‘wire’ the developing brain. This phenomenon has been widely observed in cortico-thalamic circuits, so this hypothesis is not without precedent (reviewed by [44]). Another recent *in vivo* study found that optogenetic inhibition of NRe projections to CA1 *increases* pyramidal cell activity [45], providing strong evidence to support our view of NRe’s principal overall effect on hippocampal activity is one of inhibition and not excitation, even if some sparse monosynaptic projections to pyramidal cells persist into adulthood.

While our electrophysiology data in both mouse and rat would suggest that there are no direct projections from NRe to CA1, it is possible that sparse, weak synapses could exist but, due to dendritic filtering, produce no detectable response from a somatic whole-cell patch-clamp recording. We thus carried out rabies tracing using pseudotyped SAD B19 viruses with injections made into both dorsal and ventral CA1 and also dorsal subiculum to trace inputs specifically from CA1 pyramidal cells. We failed to find evidence of a direct monosynaptic projection from NRe to CA1 pyramidal cells in either dorsal or ventral CA1. The possibility remains that the rabies virus will not retrogradely label all axon types with the same efficiency, producing a false negative. However, our control data revealed retrograde labelling of NRe neurons when the injection site included ventral subiculum, and we also found retrogradely-labelled glutamatergic neurons in regions such as entorhinal cortex. While our data cannot entirely exclude the existence of a sparse projection from NRe to CA1 pyramidal cells, if these projections do exist then they are likely to be of little functional significance.

### Circuit implications of the reuniens to hippocampus projection

*In vivo* electrophysiological studies looking at hippocampal LFPs found that only direct EC stimulation could drive a population spike in HPC, while NRe activation primarily evoked an LFP in SL-M [19,23], with the latter paper from this group arguing that low frequency stimulation of NRe likely promotes EC – HPC communication. These papers found that stimulation of both EC and NRe lead to a supralinear summation of LFP amplitude in CA1, leading the authors to reasonably conclude that both regions drive excitation within CA1.

This work and other studies have now shown that the influence of NRe over CA1 is predominantly to inhibit hippocampal activity by targeting inhibitory interneurons. Neurogliaform cells, with a predominantly NMDA receptor-driven NRe-EPSC ([22] and the present study), will likely have a much stronger influence when hippocampus is already active. Given that EC projects to both DG and CA1, and that NRe evokes strong postsynaptic responses in EC neurons, we hypothesise that much of the effect of NRe activation on increasing the magnitude of LFP in CA1 is in fact an effect of feedforward excitation from entorhinal cortex. There is one report in the literature of midline thalamic stimulation inducing population spikes in CA1 [46], but, again, this could be due to feed-forward excitation from EC.

The NRe projection to CA1, targeting neurogliaform cells, could serve to inhibit distal dendrites of selected CA1 pyramidal cell ensembles and increase the fidelity of EC – HPC communication. Another recent study, using *in vivo* juxtacellular recording in mice, found that neurogliaform cells in CA1 can decouple pyramidal cells from entorhinal-driven gamma oscillations without altering the firing rates of pyramidal cells [47]. Given that NRe projections to CA1 are highly selective for neurogliaform cells ([22,43] and the present study), we have previously suggested that the NRe recruitment of neurogliaform cells allows NRe to re-route information flow through the hippocampus, switching between memory encoding and memory retrieval [48].

Neurogliaform cells have a wide axonal arbour and broadly inhibit pyramidal cell dendrites though a combination of GABA_A_ and GABA_B_ receptor-mediated mechanisms acting via volume transmission [49], making them well placed to disengage CA1 pyramidal cells from ongoing entorhinal inputs arriving in stratum lacunosum-moleculare alongside those from nucleus reuniens. We proposed previously that the reuniens to CA1 neurogliaform projection may allow specific ensembles of pyramidal cells to be ‘disconnected’ from entorhinal cortex, creating a more permissive environment for them to be driven by CA3 inputs via Schaffer collaterals in stratum radiatum [48]. Lesions to NRe have been shown to impair consolidation, but not acquisition, of spatial memory in the Morris Water Maze in rats [50], so neurogliaform cells could play a key role in memory consolidation by supressing non-relevant activity during systems consolidation. Others have shown that NRe is essential for reconsolidation of an extinct fear memory [51–54]. Again, the NRe projection to neurogliaform cells in CA1 could play a key role in this by increasing the signal-to-noise ratio of the reactivated memory by suppressing ensembles active during similar but different episodic memories. This would be consistent with other reports of NRe being essential for the formation of long-term (but not short-term) recognition memories [55]. Neurogliaform cells have been shown to evoke a longer-lasting inhibitory current compared with other cell types such as fast-spiking basket cells [56] and acting through volume transmission [49] could make their actions too broad or slow for selecting specific ensembles, although more recent work using juxtacellular recordings in CA1 suggest that neurogliaform cell can selectively modulate network dynamics without broadly suppressing activity [47].

Prefrontal cortex projections to the hippocampus are important for impulse control [57], and lesions of NRe inhibit this function [58]. A recent suggestion for the function of the prefrontal – reuniens – hippocampal circuit is that it actively suppress ongoing memory retrieval [59], and our study provides a mechanism through which this could occur. Indeed, the large NMDA:AMPA ratio of NRe-EPSCs in neurogliaform cells that we reported here and previously [22] suggest that the inhibitory influence evoked via NRe activation is much more likely to be effective in the context of ongoing network activity. Recent behavioural data suggest the requirement for NRe in spatial memory retrieval or “online” spatial processing, but not for off- line consolidation or long-term storage [60]. Given that the largest NRe-mediated EPSCs were present in entorhinal and prefrontal cortices, it could be that NRe suppresses hippocampal activation to facilitate prefrontal – entorhinal communication during associative memory consolidation [6]. Alternatively, an inhibitory influence of NRe over hippocampus has been proposed as the mechanism through which prefrontal cortex can prevent retrieval, e.g. during extinction of fear memory [61].

Given the large NMDA receptor-mediated component of NRe-EPSCs onto neurogliaform cells, one is tempted to suggest that the role of NRe could be to increase Ca^2+^ in NGF dendrites to activate a NO-dependent suppression of inhibition [62] that could serve to enhance entorhinal-to-CA1 communication. Future behavioural experiments are required to test the function of NRe to CA1 projections, perhaps by using retrograde viruses to allow specific targeting only of those NRe neurons that project directly to CA1.

### Concluding remarks

We have found that, unique to all hippocampal or cortical regions, pyramidal cells in hippocampal region CA1 do not receive monosynaptic input from the thalamus. Even in parts of the hippocampal formation where principal cells do receive direct input from reuniens, the amplitude of the evoked currents is small and likely to be modulatory in nature. These surprise findings suggest that feedforward inhibition driven by neurogliaform cells may be main mechanism through which NRe influences hippocampal function. The apparently stronger innervation of both PFC and EC pyramidal cells by EC raises interesting questions about the flow of information through NRe (hitherto presumed to be the relaying prefrontal control over hippocampus). The existence of distinct, parallel streams of information through reuniens remains a possibility, and future research at the circuit and synaptic level should focus on NRe’s innervation of prefrontal and entorhinal regions.

## Materials and Methods

### Animals

All mouse experiments were conducted in accordance with animal protocols approved by the National Institutes of Health, or in accordance with the UK Animals (Scientific Procedures) Act 1986 after local ethical review by the Animal Welfare and Ethical Review Board at the University of Exeter under project licence PAC082CD3. We used *Nkx2-1-cre* [63]:RCE or *Nkx2-1-cre:*Ai9 and *Htr3a*-GFP [64] mice to target interneurons of MGE or CGE origin, respectively, in electrophysiological experiments. *Nkx2-1-cre* mice were obtained from Jackson laboratories (C57BL/6J-Tg(Nkx2-1-cre)2Sand/J, stock number 008661) and Htr3a- GFP mice (Tg(Htr3a-EGFP)DH30Gsat) were cryo-recovered from MMRRC (NC, USA) and back-crossed onto C57BL/6J mice (Charles River, UK). We used *Emx1*-cre mice [65] crossed with floxed TVA mice [66] to allow specific targeting of pyramidal cells for monosynaptic rabies tracing*. Emx1*-cre mice were obtained from Jackson laboratories (B6.129S2-Emx1^tm1(cre)Krj^, stock number 005628) and floxed TVA mice (LSL-R26^Tva-lacZ^) were kindly provided by Prof Dieter Saur (Technical University of Munich, Germany). All animals were maintained on a 12 h constant light / dark cycle and had access to food and water *ad libitum* and were grouped housed wherever possible. We used standard enrichment that included cardboard tubes, wooden chew blocks and nesting material.

All rat experiments were carried out in naïve male Lister Hooded rats (Envigo, UK) weighing 300 – 450 g at the start of the experiments. Animals were housed in groups of 2-4, under a 12 h light/dark cycle (light phase, 8.00 P.M. to 8.00 A.M) with *ad libitum* access to food and water. Sacrifice for *ex vivo* slices occurred 2-3 hours into the dark cycle. All animal procedures were performed in accordance with United Kingdom Animals Scientific Procedures Act (1986) and associated guidelines under project licence number PP7058522 and were approved by the University of Bristol Ethical Review Committee. All efforts were made to minimise any suffering and the number of animals used.

### Drugs and chemicals

CGP55845 (Catalogue # 1248), DNQX (Catalogue # 2312/10), D-AP5 (Catalogue # 0106/1) and picrotoxin (Catalogue # 1128) were purchased from Tocris Bioscience, and all other chemicals were purchased from Sigma-Aldrich unless otherwise stated.

### Stereotaxic injections for electrophysiology experiments: mice

For optogenetic experiments, we used *Nkx2-1*-cre:Ai9, *Nkx2-1-cre*:RCE or *Htr3a*-GFP mice of both sexes, totalling at 65 mice. with the age at the time of stereotaxic injection ranging from 2 – 7 months. Mice of both sexes were used for stereotaxic surgery. Mean weight of the mouse prior to stereotaxic surgery was 26 g (ranging from 17.5 g to 49.5 g). Two different viruses were used for stereotaxic surgeries: AAV8-hSyn-Chrimson-TdTom (UNC Viral Vector Core, USA, contributed by Ed Boyden; titre 6.3 × 10^12^ viral particles / ml). AAV8-hSyn-Chronos-GFP (UNC Viral Vector Core, USA. contributed by Ed Boyden; titre 3.1 × 10^13^ viral particles / ml).

For the surgery, the mice were anaesthetised with 5% isoflurane and anaesthesia was maintained with use of 1.5 to 2.5% isoflurane (flow rate of ∼2 Lmin^-1^ O_2_). The mice were placed on a heated pad (37 C) for the duration of the surgery and given 0.1 mg/kg of buprenorphine (buprenorphine hydrochloride, Henry Schein) subcutaneously at the start of surgery as an adjunct analgesic, plus carprofen 1 mg/kg (Rimadyl, Henry Schein) was given at a dose of 5 mg/kg subcutaneously post-surgery and on subsequent days, as required. To target nucleus reuniens, we used the following coordinates: A/P -0.8 mm, M/L 0.0 mm, D/V 3.8 mm from pia, with 300 nl of virus (infused at 100 nl min^-1^). After the surgery, the mice were allowed at least a 3-week recovery period to allow sufficient time for the expression of the viral construct. For whole cell patch clamping experiments AAV8-hSyn-Chronos-GFP or AAV8-hSyn-Chrimson-TdTom were used for *Nkx2.1*-cre:Ai9 or *Htr3a*-GFP mice, respectively, although a small number of *Htr3a-*GFP mice received AAV8-hSyn-Chronos-GFP to allow direct comparison of EPSC amplitude in the same population of neurons.

### Stereotaxic injections for electrophysiology experiments: rats

Each rat was anaesthetised with isoflurane (4% induction, 2.5-3.5% maintenance) and secured in a stereotaxic frame with the incisor bar set 3.3mm below the interaural line. Eye drops (0.1% sodium hyaluronate; Hycosan, UK) were applied and body temperature maintained at 37 °C using a homeothermic heat blanket (Harvard Apparatus, USA). The scalp was further anaesthetised using topical lidocaine (5% m/m; TEVA; UK) and disinfected with chlorhexidine, cut, and retracted. Bilateral craniotomies were made using a burr at the following coordinates with respect to Bregma: anterior-posterior (AP) – 2.0 mm, mediolateral (ML) ± 1.4 mm. Virus was injected bilaterally into nucleus reuniens via a 33-gauge 12° bevelled needle (Esslab) attached to a 5 μl Hamilton syringe which was mounted at a 10° angle in the mediolateral plane to avoid the sinus, with the eyelet of the needle facing medially. The needle was lowered 7.5 mm below the surface of the skull measured from the burr hole and 100 nl of virus was delivered via each burr hole at a rate of 200 nl/min^1^, with the needle left in situ for 10 minutes after each injection. AAV9-CaMKii-hChR2(E123T/T159C)-mCherry (Addgene 35512; 3.3 x 10^13^ genome copies/ml) obtained from University of Pennsylvania Vector Core.

### Monosynaptic retrograde tracing

For anatomical experiments we used the monosynaptic rabies tracing method that has been previously reported by others [31]. Two mouse lines (*Emx1*-cre and floxed TVA) were crossed together in order to ensure that the modified rabies virus only targets the pyramidal cells, with *Emx1-cre* mice used as controls to ensure the rabies virus did not transduce neurons in the absence of TVA. A total of 22 mice of both sexes were used, with 2 mice being excluded from the analysis due to failed injections. The age of the mice used ranged from 2 to 6 months, with pre-surgical weights from 18.9 g to 40.6 g (mean age 3.5 months, mean weight 24.6 g). To highlight the efferent projections from nucleus reuniens to hippocampus, we elected to inject into dorsal and ventral CA1 using the following coordinates: dCA1 was targeted at A/P −2 mm (relative to Bregma), M/L −1.5 mm and D/V-1.35 mm (from pia) and vCA1 at A/P −2.8 mm (relative to Bregma), M/L −2.4 mm and D/V-4.2 mm (from pia). The stereotaxic injections of AAV8-FLEX-H2B-GFP-2A-oG (Provided by John Naughton at Salk Institute Viral Vector Core, USA, titre 3.93x10^12^ viral particles / ml), followed by injection EnvA G-deleted Rabies-mCherry (Provided by John Naughton at Salk Institute Viral Vector Core, titre 6.13x10^8^ viral particles / ml; or from Viral Vector Core facility of the Kavli Institute for Systems Neuroscience, NTNU, Norway, titre 2.6 x 10^10^ viral particles / ml) 2 weeks after the initial viral injection were performed in the right hemisphere only. The mice were maintained for 2 weeks to provide optimal time for expression, and were killed by transcardial perfusion / fixation with 4% paraformaldehyde (Cat number P6148 Sigma-Aldrich, UK) in 0.1 M phosphate buffer.

Following the transcardial perfusion, the brains were dissected out and post-fixed for 24 h in 4% pfa solution, after which they were cryoprotected using the 30% sucrose in PBS solution (Catalogue # P4417). Once cryoprotected, the brains were sliced at 50 microns using the freezing microtome (Leica, SM2010 R). Selected slices (1 in 5 serially, increasing to 1 in 3 between -0.5 and −1.8 Bregma to ensure thorough representation of nucleus Reuniens of the thalamus across the anterior-posterior axes) were mounted using the Hard Set mounting medium with DAPI (Vectashield, Vector Lab, Catalogue # H-1500-10) and the fluorescent fibres were visualised with CoolLED on Nikon Eclipse E800, using 4x objective. Representative photos of projections patterns can be found on figure 1 – figure supplement 1.

### Slice preparation and electrophysiology

Mice of age two months and above were used for stereotaxic surgeries. A minimum of 3 weeks recovery period following the stereotaxic surgery was allowed. Mice were anesthetised with isoflurane and the brain was rapidly dissected out in room temperature NMDG cutting solution, containing (in mM): 135 NMDG (Catalogue # M2004), 20 Choline bicarbonate (Catalogue # C7519), 10 glucose (Catalogue # G7021), 1.5 MgCl_2_ (Catalogue # M9297), 1.2 KH_2_PO_4_ (Catalogue # P0662), 1 KCl (Catalogue # P5405), 0.5 CaCl_2_ (Catalogue # C5670), saturated with 95% O_2_ and 5% CO_2_ (pH 7.3-7.4). Coronal or horizontal slices (400 um) were cut to target prefrontal cortex and dorsal CA1 vs ventral CA1, subiculum and entorhinal cortex, respectively, using a VT-1200S vibratome (Leica Microsystems, Germany). Afterwards, the slices were transferred into a chamber containing recording aCSF, composed of (in mM): 130 NaCl (Catalogue # S5886), 24 NaHCO_3_ (Catalogue # S6014), 3.5 KCl, 1.25 NaH_2_PO_4_, 2.5 CaCl_2_, 1.5 MgCl_2_, and 10 glucose, saturated with 95% O_2_ and 5% CO_2_ (pH 7.4) and placed in a water bath at 37 °C for 30 minutes, following which they were kept at room temperature until recording.

For rat experiments, after a minimum of 10 days following viral injection, animals were anaesthetised with 4% isoflurane and decapitated. Brains were rapidly removed and placed into ice-cold sucrose solution (in mM: 189 sucrose, 26 NaHCO_3_, 10 D-glucose, 5 MgSO_4_, 3 KCl, 1.25 NaH_2_PO_4_, 0.2 CaCl_2_) bubbled with 95 % O_2_/5% CO_2_. Dorsal hippocampal recordings were made from parasagittal or coronal slices and ventral hippocampus from horizontal slices, cut at 350 μm thickness using a vibratome (7000smz-2, Camden Instruments), before incubation at 34 °C for 1 hour after dissection in a slice holding chamber filled with artificial cerebrospinal fluid (aCSF, in mM: 124 NaCl, 26 NaHCO_3_, 10 D-glucose, 3 KCl, 2 CaCl_2_, 1.25 NaH2PO_4_, 1 MgSO_4_). Slices were subsequently stored at room temperature until use.

For recordings individual mouse brain slices were attached onto 0.1% poly-L-lysine (Sigma Aldrich, Catalogue # P8920) coated glass slides and placed into the upright microscope and visualized using infrared differential interference contrast microscopy (Olympus BX51 or Scientifica SliceScope). CoolLED pE-4000 system was used to visualise the fibres as well as interneurons, and to provide optogenetic stimulation. The slices were submerged in recording aCSF, warmed to 32-34 °C, and the rate of perfusion was kept at 5ml/min. The recording electrodes were typically 3-5 MW size and were pulled from borosilicate glass (World Precision Instruments). The intracellular solution used had the following composition (in mM): 135 Cs-methanesulfonate (Catalogue # C1426), 8 NaCl, 10 HEPES (Catalogue # H3375), 0.5 EGTA (Catalogue # E3889), 4 MgATP (Catalogue # A9187), 0.3 Na-GTP Catalogue # (G8877), 5 QX314 (Catalogue # 552233), plus 2 mg/ml biocytin (VWR International, UK), at pH 7.25 adjusted with CsOH and 285 mOsm.

For rat electrophysiological recordings, slices at were placed in a submerged recording chamber and perfused with 34°C aCSF at ∼2ml/min^1^. A stimulating electrode (FH-Co, CBABAP50, USA) was placed in stratum lacunosum moleculare. Recordings were made from neurons in stratum pyramidale or lacunosum moleculare, targetted under infra-red illumination with an upright microscope and whole-cell patch clamped using 2-6 MΩ boroscillicate glass electrodes (GC150-10F, Harvard Apparatus) filled with potassium gluconate internal (in mM: 120 k-gluconate, 40 HEPES, 10 KCl, 2 NaCl, 2 MgATP, 1 MgCl, 0.3 NaGTP, 0.2 EGTA and either 0.1 Alexa-594 hydrazide or 2.7 biocytin, pH 7.25, 285 mOsm). Recordings were obtained using a Molecular Devices Multiclamp 700B, filtered at 4 KHz and digitized at a sample frequency ≥20 KHz with WinLTP2.30 (Anderson & Collingridge 2007) or pClamp10 software. Resting membrane potential (RMP) was recorded immediately after entering the whole-cell configuration, thereafter neurons were kept at −70 mV by injection of constant current. Intrinsic membrane properties were recorded by injection of square-wave hyperpolarising and depolarising currents. Optogenetic stimulation was applied with a 470 nm LED (M470L3, Thorlabs) triggered by 2ms TTL pulses sent to an LEDD1B driver (Thorlabs) via a 40x immersion objective (Olympus or Nikon). In all experiments transduction of nucleus reuniens was confirmed by recording of photosensitive cells in hippocampal and/or medial prefrontal cortex slices [30]. Cell morphology was confirmed using either fluorescent visualisation of Alexa594 signal, or post-hoc biocytin staining.

A train stimulation with 5 pulses of 470 nm or 660 nm was used to excite the Chronos [67] or Chrimson opsins, respectively. The presence or absence of responses was recorded in voltage clamp mode. Cells that were found to have a response to a train stimulation were then switched onto repeated single pulse protocol (ISI of 10 s), and the AMPA response was recorded at a holding potential of −70 mV. GABA-R antagonists were bath applied from the start in hippocampus and subiculum, but not in EC or PFC due to epileptiform activity being observed upon NRe stimulation with GABA-R antagonists present. The extracellular GABA_A_ and GABA_B_ receptor antagonists used were picrotoxin (100 µM) and CGP55845 (1 µM). 10 µm of DNQX was added to abolish the AMPA current at −70 mV, after which the cell was switched to +40 mV to record the NMDA current. To confirm the identity of NMDA current, D- AP 5 (100 µM) was added at the end of the recording. Whole-cell patch-clamp recordings were made using a Multiclamp 700A or 700B amplifier (Molecular Devices, Sunnyvale, CA). Signals were filtered at 3 kHz and digitized at 10 kHz using a Digidata 1322A or 1440A and pClamp 9.2 or 10.2 (Molecular Devices, USA). Recordings were not corrected for a liquid junction potential. The recordings were then imported into IgorPro (Wavemetrics, OR) using Neuromatic (Thinkrandom, UK) for further analysis.

Total number and age range of mice (on experimental day) used for whole cell patch clamping data was: 28 mice of both sexes aged 4-7 months old for hippocampal formation (Figures 1 & 2), 15 mice of both sexes aged 4-7 months old for entorhinal cortex (Figure 5) and 21 mice of both sexes aged 4-8 months old for prefrontal cortex (Figure 4).

### Post hoc morphological recovery

Mouse tissue slices were post-fixed in 4% PFA solution for an hour after patch clamping recordings and then transferred to PBS (Melford, UK). Slices were washed thrice in 0.1 M PBS solution, followed by three washes in PBS with 0.5% Triton X (Sigma, UK). Two blocking steps were used: 100mM of Glycine (Sigma, UK) incubation for 20 mins and then 1-hour long incubation with blocking buffer with 5% goat serum (VectorLabs, UK) at room temperature. Incubation with streptavidin (1:500 dilution, VectorLabs, UK) was done for 2 hours at RT in a carrier solution consisting of 5% goat serum and PBS. Finally, the slices were washed in PBS and transferred to 30% sucrose (Thermofisher, UK) until cryoprotected. Then slices were re-sectioned at 100-micron thickness using SM2010R freezing microtome (Leica, UK)) at −20°C and mounted onto glass-slides and HardSet mounting medium with DAPI was used (VectorLabs, UK).

For rat experiments, following recording, slices were fixed in 4% paraformaldehyde overnight and subsequently stored in 0.1 M phosphate buffer (PB) until staining. Slices were washed 6 x 10 min in PB, then incubated in 3% H_2_O_2_ in PB for 30 min to block endogenous peroxidase activity, then washed again as above before incubation in 1% (vol/vol) avidin–biotinylated HRP complex (ABC) in PB containing 0.1% (vol/vol) Triton X-100 at RT for 3 hrs. Slices were then washed as above before incubation in DAB solution (1x gold + 1x silver tablet in 15 mL distilled water) for 5-10 min until staining of neuronal structures was visible. The reaction was stopped by transferring slices to cold PB, followed by further washing as above. Slices were mounted, cover slipped using Mowiol mounting media and allowed to dry overnight before imaging.

### Data Analysis

For quality control, cells with changes in input resistance of over 20% throughout the course of the experiment were excluded from the data analysis. The AMPA receptor-mediated EPSC was determined as the maximal EPSC peak at −70mV and NMDA receptor-mediated EPSC as the highest peak at +40 mV. GraphPad Prism (GraphPad, CA) was used for statistical analysis. Data were tested for normality using the D’Agostino and Pearson test and subsequently analysed by parametric or nonparametric tests as appropriate. Unless otherwise stated, all values are mean ± SEM. RNAseq data from [32] were downloaded from NCBI Gene Expression omnibus [68], accession GSE67403) and analysed in R Statistical Software v4.3.1 [69] and RStudio v 2023.6.1.524 [70], using the tidyverse [71] and ggplot2 packages [72].

For rat experiments, traces from intrinsic electrophysiological experiments were imported into MATLAB and analysed using custom written code. Synaptic responses were analysed using MATLAB or WinLTP software [73] .

## Acknowledgements

This work was supported by Biotechnology and Biological Sciences Research Council grant BB/P001475/1 (MTC), an NIH intramural award (CJM), a Wellcome Trust Joint Investigator Award 206401/Z/17/Z (ZIB), and Biotechnology and Biological Sciences Research Council grants BB/X000915/1 and BB/L001896/1 (ZIB, PJB). Salary support from SK was provided by Alzheimer’s Research UK Interdisciplinary research grant ARUK-IRG2017B-4 (MTC). GMS and ESB are both GW4 BioMed doctoral training program students funded by the Medical Research Council (MR/N0137941/1). We are grateful to Prof John Aggleton (University of Cardiff) for advice and discussions on anatomy. We gratefully acknowledge the support of the Inger and George Simpson Biological Psychiatry Scholarships. For the purpose of open access, the authors have applied a Creative Commons Attribution (CC-BY) licence to any Author Accepted Manuscript version arising from this submission.

## References

1. Preston AR, Eichenbaum H. Interplay of Hippocampus and Prefrontal Cortex in Memory. Curr Biol. 2013;23: R764–R773. doi:10.1016/j.cub.2013.05.041

2. Euston DR, Gruber AJ, McNaughton BL. The Role of Medial Prefrontal Cortex in Memory and Decision Making. Neuron. 2012;76: 1057–1070. doi:10.1016/j.neuron.2012.12.002

3. Dolleman-van der Weel MJ, Witter MP. THE THALAMIC MIDLINE NUCLEUS REUNIENS: POTENTIAL RELEVANCE FOR SCHIZOPHRENIA AND EPILEPSY. Neurosci Biobehav Rev. 2020;119: 422–439. doi:10.1016/j.neubiorev.2020.09.033

4. Jones MW, Wilson MA. Theta Rhythms Coordinate Hippocampal–Prefrontal Interactions in a Spatial Memory Task. Plos Biol. 2005;3: e402. doi:10.1371/journal.pbio.0030402

5. Tang W, Shin JD, Jadhav SP. Multiple time-scales of decision making in the hippocampus and prefrontal cortex. Elife. 2021;10: e66227. doi:10.7554/elife.66227

6. Takehara-Nishiuchi K, Maal-Bared G, Morrissey MD. Increased Entorhinal–Prefrontal Theta Synchronization Parallels Decreased Entorhinal–Hippocampal Theta Synchronization during Learning and Consolidation of Associative Memory. Front Behav Neurosci. 2012;5: 90. doi:10.3389/fnbeh.2011.00090

7. Hartley T, Lever C, Burgess N, O’Keefe J. Space in the brain: how the hippocampal formation supports spatial cognition. Philosophical Transactions Royal Soc B Biological Sci. 2014;369: 20120510. doi:10.1098/rstb.2012.0510

8. Andrianova L, Yanakieva S, Margetts-Smith G, Kohli S, Brady ES, Aggleton JP, et al. No evidence from complementary data sources of a direct glutamatergic projection from the mouse anterior cingulate area to the hippocampal formation. eLife. 2023;12. doi:10.7554/elife.77364

9. Mathiasen ML, O’Mara SM, Aggleton JP. The anterior thalamic nuclei and nucleus reuniens: So similar but so different. Neurosci Biobehav Rev. 2020;119: 268–280. doi:10.1016/j.neubiorev.2020.10.006

10. Griffin AL. Role of the thalamic nucleus reuniens in mediating interactions between the hippocampus and medial prefrontal cortex during spatial working memory. Frontiers Syst Neurosci. 2015;9: 29. doi:10.3389/fnsys.2015.00029

11. Dolleman-van der Weel MJ, Griffin AL, Ito HT, Shapiro ML, Witter MP, Vertes RP, et al. The nucleus reuniens of the thalamus sits at the nexus of a hippocampus and medial prefrontal cortex circuit enabling memory and behavior. Learn Memory. 2019;26: 191–205. doi:10.1101/lm.048389.118

12. Vertes RP, Hoover WB, Szigeti-Buck K, Leranth C. Nucleus reuniens of the midline thalamus: Link between the medial prefrontal cortex and the hippocampus. Brain Res Bull. 2007;71: 601–609. doi:10.1016/j.brainresbull.2006.12.002

13. Dolleman-van der Weel MJ, Morris RGM, Witter MP. Neurotoxic lesions of the thalamic reuniens or mediodorsal nucleus in rats affect non-mnemonic aspects of watermaze learning. Brain Struct Funct. 2009;213: 329–342. doi:10.1007/s00429-008-0200-6

14. Herkenham M. The connections of the nucleus reuniens thalami: Evidence for a direct thalamo-hippocampal pathway in the rat. J Comp Neurol. 1978;177: 589–609. doi:10.1002/cne.901770405

15. Wouterlood FG, Saldana E, Witter MP. Projection from the nucleus reuniens thalami to the hippocampal region: Light and electron microscopic tracing study in the rat with the anterograde tracer Phaseolus vulgaris-leucoagglutinin. J Comp Neurol. 1990;296: 179–203. doi:10.1002/cne.902960202

16. Schlecht M, Jayachandran M, Rasch GE, Allen TA. Dual projecting cells linking thalamic and cortical communication routes between the medial prefrontal cortex and hippocampus. Neurobiol Learn Mem. 2022;188: 107586. doi:10.1016/j.nlm.2022.107586

17. Wouterlood FG. Innervation of Entorhinal Principal Cells by Neurons of the Nucleus Reuniens Thalami. Anterograde PHA-L Tracing Combined with Retrograde Fluorescent Tracing and Intracellular Injection with Lucifer Yellow in the Rat. Eur J Neurosci. 1991;3: 641–647. doi:10.1111/j.1460-9568.1991.tb00850.x

18. McKenna JT, Vertes RP. Afferent projections to nucleus reuniens of the thalamus. J Comp Neurol. 2004;480: 115–142. doi:10.1002/cne.20342

19. Dolleman-van der Weel MJ, Silva FHL da, Witter MP. Nucleus Reuniens Thalami Modulates Activity in Hippocampal Field CA1 through Excitatory and Inhibitory Mechanisms. J Neurosci. 1997;17: 5640–5650. doi:10.1523/jneurosci.17-14-05640.1997

20. Ito HT, Zhang S-J, Witter MP, Moser EI, Moser M-B. A prefrontal–thalamo– hippocampal circuit for goal-directed spatial navigation. Nature. 2015;522: 50–55. doi:10.1038/nature14396

21. Duan AR, Varela C, Zhang Y, Shen Y, Xiong L, Wilson MA, et al. Delta Frequency Optogenetic Stimulation of the Thalamic Nucleus Reuniens Is Sufficient to Produce Working Memory Deficits: Relevance to Schizophrenia. Biol Psychiat. 2015;77: 1098–1107. doi:10.1016/j.biopsych.2015.01.020

22. Chittajallu R, Wester JC, Craig MT, Barksdale E, Yuan XQ, Akgül G, et al. Afferent specific role of NMDA receptors for the circuit integration of hippocampal neurogliaform cells. Nat Commun. 2017;8: 152. doi:10.1038/s41467-017-00218-y

23. Dolleman-van der Weel MJ, Silva FHL da, Witter MP. Interaction of nucleus reuniens and entorhinal cortex projections in hippocampal field CA1 of the rat. Brain Struct Funct. 2017;222: 2421–2438. doi:10.1007/s00429-016-1350-6

24. Klapoetke NC, Murata Y, Kim SS, Pulver SR, Birdsey-Benson A, Cho YK, et al. Independent optical excitation of distinct neural populations. Nat Methods. 2014;11: 338–346. doi:10.1038/nmeth.2836

25. Lorente De Nó R. Studies on the Structure of the Cerebral Cortex II. Continuation of the Study of the Ammonic System. J Psychol Neurol. 1934;46: 113–177.

26. Ding S. Comparative anatomy of the prosubiculum, subiculum, presubiculum, postsubiculum, and parasubiculum in human, monkey, and rodent. J Comp Neurol. 2013;521: 4145–4162. doi:10.1002/cne.23416

27. Canto CB, Witter MP. Cellular properties of principal neurons in the rat entorhinal cortex. II. The medial entorhinal cortex Hippocampus. 2012;22: 1277–1299. doi:10.1002/hipo.20993

28. Canto CB, Witter MP. Cellular properties of principal neurons in the rat entorhinal cortex. I. The lateral entorhinal cortex Hippocampus. 2012;22: 1256–1276. doi:10.1002/hipo.20997

29. Pastoll H, Garden DL, Papastathopoulos I, Sürmeli G, Nolan MF. Inter- and intra-animal variation in the integrative properties of stellate cells in the medial entorhinal cortex. eLife. 2020;9: e52258. doi:10.7554/elife.52258

30. Banks PJ, Warburton EC, Bashir ZI. Plasticity in Prefrontal Cortex Induced by Coordinated Synaptic Transmission Arising from Reuniens/Rhomboid Nuclei and Hippocampus. Cereb Cortex Commun. 2021;2: tgab029. doi:10.1093/texcom/tgab029

31. Sun Y, Nguyen AQ, Nguyen JP, Le L, Saur D, Choi J, et al. Cell-Type-Specific Circuit Connectivity of Hippocampal CA1 Revealed through Cre-Dependent Rabies Tracing. Cell Reports. 2014;7: 269–280. doi:10.1016/j.celrep.2014.02.030

32. Cembrowski MS, Bachman JL, Wang L, Sugino K, Shields BC, Spruston N. Spatial Gene-Expression Gradients Underlie Prominent Heterogeneity of CA1 Pyramidal Neurons. Neuron. 2016;89: 351–368. doi:10.1016/j.neuron.2015.12.013

33. Ding S-L, Yao Z, Hirokawa KE, Nguyen TN, Graybuck LT, Fong O, et al. Distinct Transcriptomic Cell Types and Neural Circuits of the Subiculum and Prosubiculum along the Dorsal-Ventral Axis. Cell Reports. 2020;31: 107648. doi:10.1016/j.celrep.2020.107648

34. Wee RWS, MacAskill AF. Biased Connectivity of Brain-wide Inputs to Ventral Subiculum Output Neurons. Cell Reports. 2020;30: 3644–3654.e6. doi:10.1016/j.celrep.2020.02.093

35. Andrianova L, Brady ES, Margetts-Smith G, Kohli S, Cavanagh J, McBain CJ, et al. Hippocampus does not appear to be a major target of thalamic nucleus reuniens. bioRxiv. 2021; 2021.09.30.462517. doi:10.1101/2021.09.30.462517

36. Goswamee P, Leggett E, McQuiston AR. Nucleus Reuniens Afferents in Hippocampus Modulate CA1 Network Function via Monosynaptic Excitation and Polysynaptic Inhibition. Front Cell Neurosci. 2021;15: 660897. doi:10.3389/fncel.2021.660897

37. Bloss EB, Cembrowski MS, Karsh B, Colonell J, Fetter RD, Spruston N. Single excitatory axons form clustered synapses onto CA1 pyramidal cell dendrites. Nat Neurosci. 2018;21: 353–363. doi:10.1038/s41593-018-0084-6

38. Foggetti A, Baccini G, Arnold P, Schiffelholz T, Wulff P. Spiny and Non-spiny Parvalbumin-Positive Hippocampal Interneurons Show Different Plastic Properties. Cell Rep. 2019;27: 3725–3732.e5. doi:10.1016/j.celrep.2019.05.098

39. Ziółkowska M, Sotoudeh N, Cały A, Puchalska M, Pagano R, Śliwińska MA, et al. Projections from thalamic nucleus reuniens to hippocampal CA1 area participate in context fear extinction by affecting extinction-induced molecular remodeling of excitatory synapses. eLife. 2025;13: RP101736. doi:10.7554/elife.101736

40. Zhang W, Vazquez L, Apperson M, Kennedy MB. Citron Binds to PSD-95 at Glutamatergic Synapses on Inhibitory Neurons in the Hippocampus. J Neurosci. 1999;19: 96–108. doi:10.1523/jneurosci.19-01-00096.1999

41. Graf ER, Zhang X, Jin S-X, Linhoff MW, Craig AM. Neurexins Induce Differentiation of GABA and Glutamate Postsynaptic Specializations via Neuroligins. Cell. 2004;119: 1013– 1026. doi:10.1016/j.cell.2004.11.035

42. Pelkey KA, Barksdale E, Craig MT, Yuan X, Sukumaran M, Vargish GA, et al. Pentraxins Coordinate Excitatory Synapse Maturation and Circuit Integration of Parvalbumin Interneurons. Neuron. 2015;85: 1257–1272. doi:10.1016/j.neuron.2015.02.020

43. Leprince E, Dard RF, Mortet S, Filippi C, Giorgi-Kurz M, Bourboulou R, et al. Extrinsic control of the early postnatal CA1 hippocampal circuits. Neuron. 2023. doi:10.1016/j.neuron.2022.12.013

44. Molnár Z, Luhmann HJ, Kanold PO. Transient cortical circuits match spontaneous and sensory-driven activity during development. Science. 2020;370: eabb2153. doi:10.1126/science.abb2153

45. Torromino G, Loffredo V, Cavezza D, Sonsini G, Esposito F, Crevenna AH, et al. Thalamo-hippocampal pathway regulates incidental memory load in mice. Biorxiv. 2022; 2021.08.07.453742. doi:10.1101/2021.08.07.453742

46. Bertram EH, Zhang DX. Thalamic excitation of hippocampal CA1 neurons: a comparison with the effects of CA3 stimulation. Neuroscience. 1999;92: 15–26. doi:10.1016/s0306-4522(98)00712-x

47. Sakalar E, Klausberger T, Lasztóczi B. Neurogliaform cells dynamically decouple neuronal synchrony between brain areas. Science. 2022;377: 324–328. doi:10.1126/science.abo3355

48. Craig MT, Witton J. A cellular switchboard in memory circuits. Science. 2022;377: 262–263. doi:10.1126/science.add2681

49. Oláh S, Füle M, Komlósi G, Varga C, Báldi R, Barzó P, et al. Regulation of cortical microcircuits by unitary GABA-mediated volume transmission. Nature. 2009;461: 1278– 1281. doi:10.1038/nature08503

50. Loureiro M, Cholvin T, Lopez J, Merienne N, Latreche A, Cosquer B, et al. The Ventral Midline Thalamus (Reuniens and Rhomboid Nuclei) Contributes to the Persistence of Spatial Memory in Rats. J Neurosci. 2012;32: 9947–9959. doi:10.1523/jneurosci.0410-12.2012

51. Sierra RO, Pedraza LK, Zanona QK, Santana F, Boos FZ, Crestani AP, et al. Reconsolidation-induced rescue of a remote fear memory blocked by an early cortical inhibition: Involvement of the anterior cingulate cortex and the mediation by the thalamic nucleus reuniens. Hippocampus. 2017;27: 596–607. doi:10.1002/hipo.22715

52. Troyner F, Bicca MA, Bertoglio LJ. Nucleus reuniens of the thalamus controls fear memory intensity, specificity and long-term maintenance during consolidation. Hippocampus. 2018;28: 602–616. doi:10.1002/hipo.22964

53. Troyner F, Bertoglio LJ. Nucleus reuniens of the thalamus controls fear memory reconsolidation. Neurobiol Learn Mem. 2021;177: 107343. doi:10.1016/j.nlm.2020.107343

54. Troyner F, Bertoglio LJ. Thalamic nucleus reuniens regulates fear memory destabilization upon retrieval. Neurobiol Learn Mem. 2020;175: 107313. doi:10.1016/j.nlm.2020.107313

55. Barker GRI, Warburton EC. A Critical Role for the Nucleus Reuniens in Long-Term, But Not Short-Term Associative Recognition Memory Formation. J Neurosci. 2018;38: 3208– 3217. doi:10.1523/jneurosci.1802-17.2017

56. Szabadics J, Tamás G, Soltesz I. Different transmitter transients underlie presynaptic cell type specificity of GABAA,slow and GABAA,fast. Proc Natl Acad Sci. 2007;104: 14831– 14836. doi:10.1073/pnas.0707204104

57. Chudasama Y, Doobay VM, Liu Y. Hippocampal-Prefrontal Cortical Circuit Mediates Inhibitory Response Control in the Rat. J Neurosci. 2012;32: 10915–10924. doi:10.1523/jneurosci.1463-12.2012

58. Prasad JA, Macgregor EM, Chudasama Y. Lesions of the thalamic reuniens cause impulsive but not compulsive responses. Brain Struct Funct. 2013;218: 85–96. doi:10.1007/s00429-012-0378-5

59. Anderson MC, Bunce JG, Barbas H. Prefrontal–hippocampal pathways underlying inhibitory control over memory. Neurobiol Learn Mem. 2016;134: 145–161. doi:10.1016/j.nlm.2015.11.008

60. Mei H, Logothetis NK, Eschenko O. The activity of thalamic nucleus reuniens is critical for memory retrieval, but not essential for the early phase of “off-line” consolidation. Learn Memory. 2018;25: 129–137. doi:10.1101/lm.047134.117

61. Anderson MC, Floresco SB. Prefrontal-hippocampal interactions supporting the extinction of emotional memories: the retrieval stopping model. Neuropsychopharmacol. 2021; 1–16. doi:10.1038/s41386-021-01131-1

62. Li G, Stewart R, Canepari M, Capogna M. Firing of Hippocampal Neurogliaform Cells Induces Suppression of Synaptic Inhibition. J Neurosci. 2014;34: 1280–1292. doi:10.1523/jneurosci.3046-13.2014

63. Xu Q, Tam M, Anderson SA. Fate mapping Nkx2.1-lineage cells in the mouse telencephalon. J Comp Neurol. 2008;506: 16–29. doi:10.1002/cne.21529

64. Gong S, Zheng C, Doughty ML, Losos K, Didkovsky N, Schambra UB, et al. A gene expression atlas of the central nervous system based on bacterial artificial chromosomes. Nature. 2003;425: 917–925. doi:10.1038/nature02033

65. Guo H, Hong S, Jin X-L, Chen R-S, Avasthi PP, Tu Y-T, et al. Specificity and Efficiency of Cre-Mediated Recombination in Emx1–cre Knock-in Mice. Biochem Bioph Res Co. 2000;273: 661–665. doi:10.1006/bbrc.2000.2870

66. Seidler B, Schmidt A, Mayr U, Nakhai H, Schmid RM, Schneider G, et al. A Cre-loxP-based mouse model for conditional somatic gene expression and knockdown in vivo by using avian retroviral vectors. Proc National Acad Sci. 2008;105: 10137–10142. doi:10.1073/pnas.0800487105

67. Klapoetke NC, Murata Y, Kim SS, Pulver SR, Birdsey-Benson A, Cho YK, et al. Independent optical excitation of distinct neural populations. Nat Methods. 2014;11: 338–346. doi:10.1038/nmeth.2836

68. Edgar R, Domrachev M, Lash AE. Gene Expression Omnibus: NCBI gene expression and hybridization array data repository. Nucleic Acids Res. 2002;30: 207–210. doi:10.1093/nar/30.1.207

69. Team RC. R: A Language and Environment for Statistical Computing. 2021. Available: https://www.R-project.org/

70. team P. RStudio: Integrated Development Environment for R. 2023. Available: http://www.posit.co/

71. Wickham H, Averick M, Bryan J, Chang W, McGowan LD, François R, et al. Welcome to the tidyverse. Journal of Open Source Software. 2019;4: 1686. doi:10.21105/joss.01686

72. Wickham H. ggplot2, Elegant Graphics for Data Analysis. 2016; 11–31. doi:10.1007/978-3-319-24277-4_2

73. Anderson WW, Collingridge GL. Capabilities of the WinLTP data acquisition program extending beyond basic LTP experimental functions. J Neurosci Methods. 2007;162: 346–356. doi:10.1016/j.jneumeth.2006.12.018

