## Supplementary figures for "Thalamic nucleus reuniens preferentially targets inhibitory interneurons over pyramidal cells in hippocampal CA1 region"

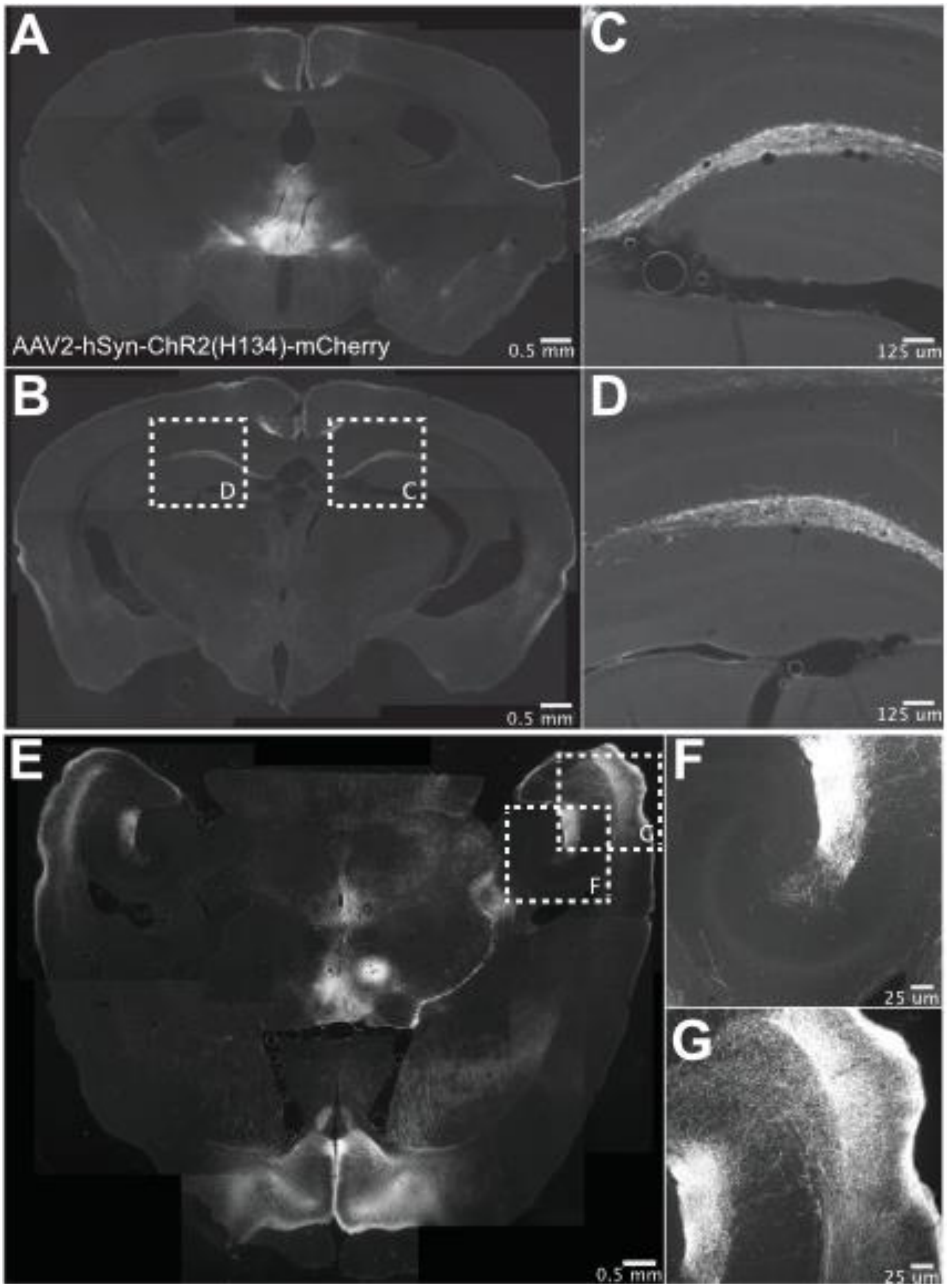


**Figure S1: NRe axon distribution in coronal and horizontal sections. A**, coronal section, showing injection site, **B,** more caudal section, showing the fibres in dorsal hippocampus, localised to S-LM. **C,** and **D,** show hatched areas from **B,** on a higher magnification.

**E,** horizontal slice, showing dense projections from nucleus reuniens in ventral prefrontal cortex, ventral CA1 and subiculum (**F**) and entorhinal cortex (**G**).

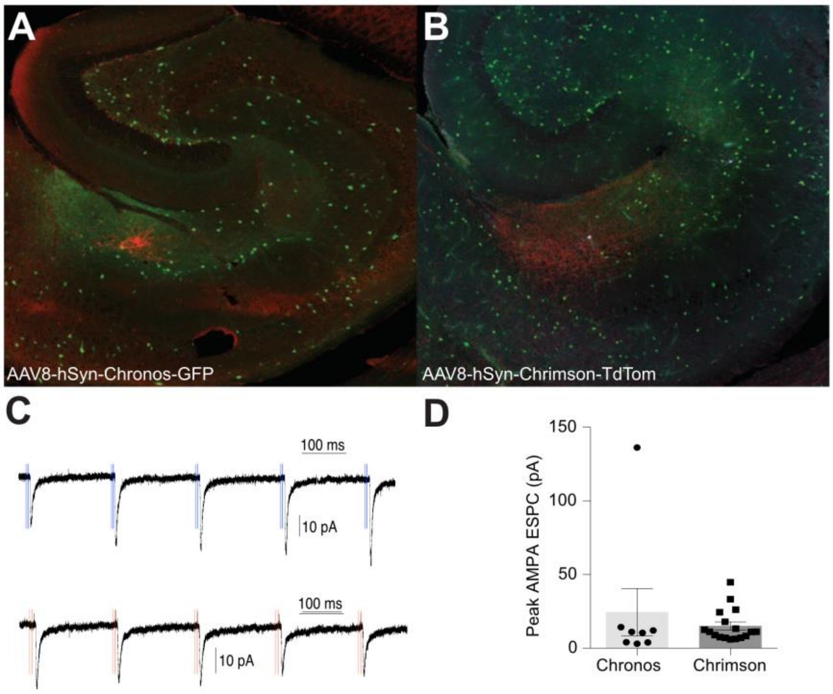


**Figure S2: No difference in NRe-EPSC between Chronos and Chrimson, in CGE-derived neurogliaform cells in CA1. A,** representative image of Chronos-GFP NRe fibres (green) in a *Htr3a*-GFP mouse. **B**, representative image of Chrimson-TdTom NRe fibres (red) in a *Htr3a*-GFP mouse. **C,** representative traces for NRe-EPSCs (AMPA-mediated) in CA1 CGE-derived neurogliaform cells from NRe axons transduced with either AAV8-hSyn-Chronos-GFP (upper trace) or AAV8-hSyn-Chrimson-TdTom (lower trace). **D**, no significant difference in median peak AMPA-mediated NRe-EPSC current between fibres expressing Chronos or Chrimson (Chronos *vs* Chrimson: 10.8 (IQR: 4.0 to 13.7, n=8) *vs* 11.2 (IQR: 7.3 to 21.2, n=17); p=0.4143, Mann-Whitney test).

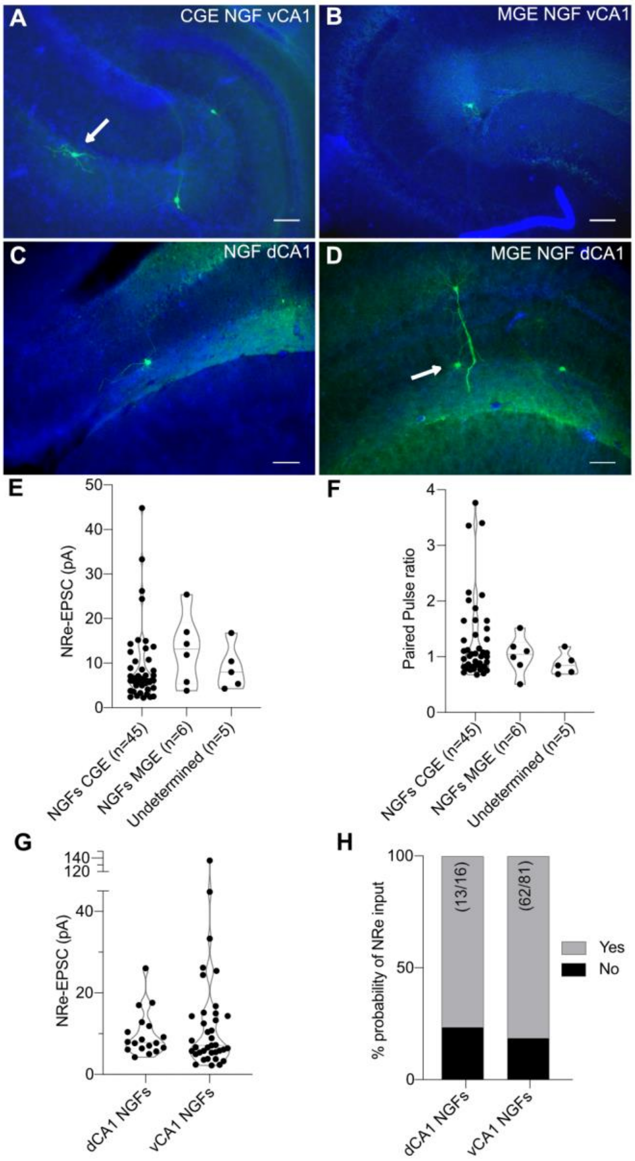


**Figure S3: comparison of NRe input to MGE vs CGE-born neurogliaform cells, and between dCA1 and vCA1.** Representative images of **A**, CGE-derived NGF in vCA1 (arrow); **B**, MGE-derived NGF in vCA1; **C**, putative CGE-derived NGF in dCA1 (TdTom -ve in Nkx2.1-cre:Ai9 mouse); **D**, MGE-derived in dCA1 (arrow). **E,** AMPA-R mediated NRe did not vary significantly between NGFs of different embryonic origin (CGE *vs* MGE *vs* indeterminate origin: 9.12 $\pm$ 1.25 pA *vs* 13.0 $\pm$ 3.2pA *vs* 8.94 $\pm$ 2.23pA; p=0.34, Kruskal-Wallis test). Similarly, **F**, NRe-EPSC paired pulse ratio did not vary by embryonic origin (CGE *vs* mGE *vs* indeterminate origin: 1.27 $\pm$ 0.11 *vs vs* 1.03$\pm$0.14 *vs* 0.88 $\pm$ 0.09; p=0.39, Kruskal-Wallis test). **G,** mean AMPA NRe-EPSC did not significantly differ between dCA1 and vCA1 NGFs, pooled across embryonic origin (dCA1 *vs* vCA1 9.88 $\pm$ 1.3 pA *vs* 14.3 $\pm$ 3.7pA; p=0.6504, Mann Whitney test). **H**, probability of NRe input in neurogliaform cells in dorsal and ventral CA1 was 76.5% and 81.3%, respectively.

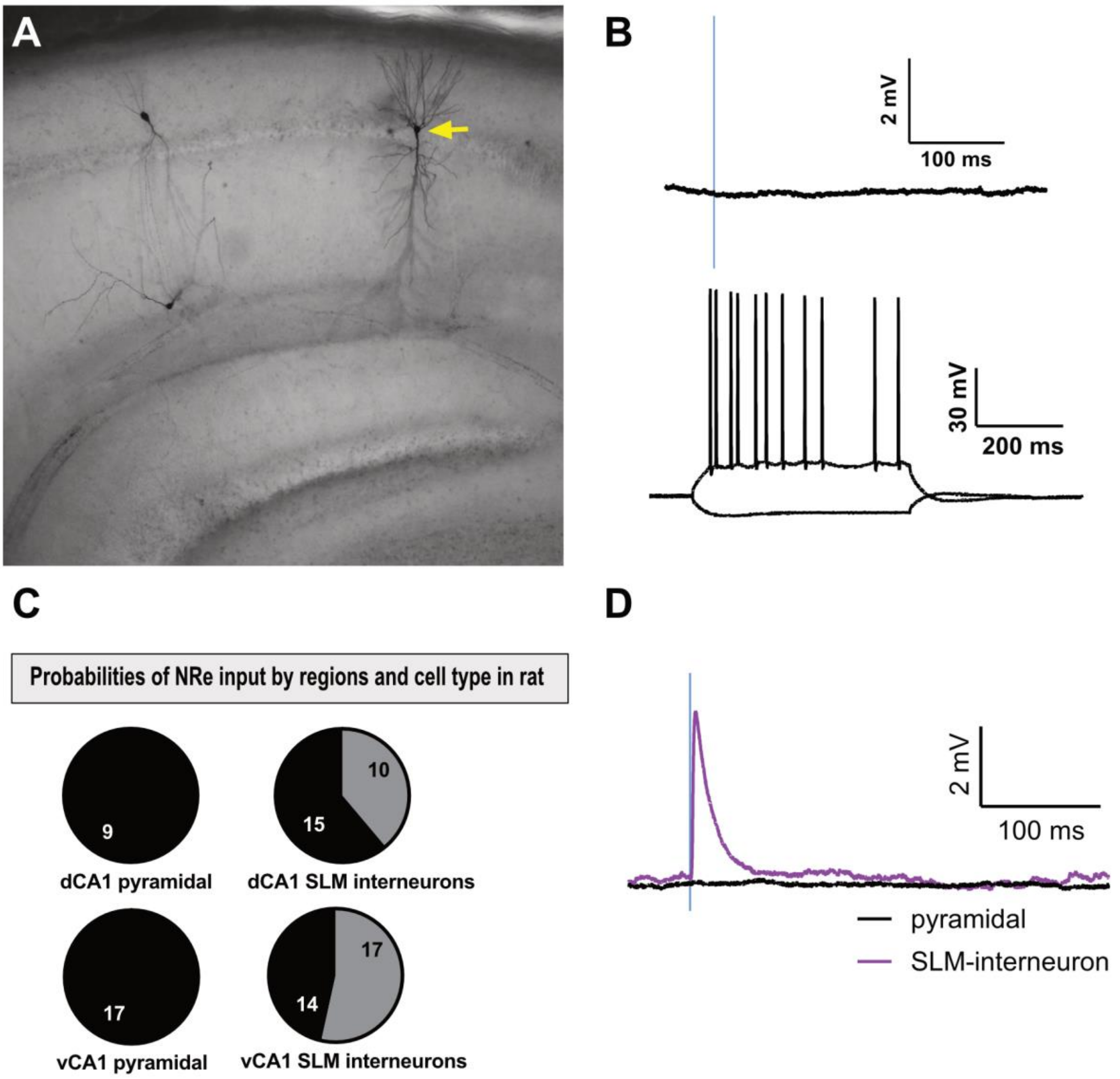


**Figure S4: Optogenetic activation of NRe axons in rat hippocampus evokes a post-synaptic response in GABAergic interneurons but not pyramidal cells in either dorsal or ventral CA1. A**, *post hoc* recovery of CA1 neurons in dorsal hippocampus of rat. **B,** lack of EPSP for pyramidal cell shown in A in response to optogenetic stimulation (top), plus response to depolarising and hyperpolarising current steps (bottom). **C**, total connection probabilities for CA1 pyramidal cells and GABAergic neurons in SL-M, with positive responses in grey and null responses in black. Data from 21 rats. **D,** example EPSP from an SLM-interneuron (purple) and a pyramidal cell (black) recorded from ventral CA1.

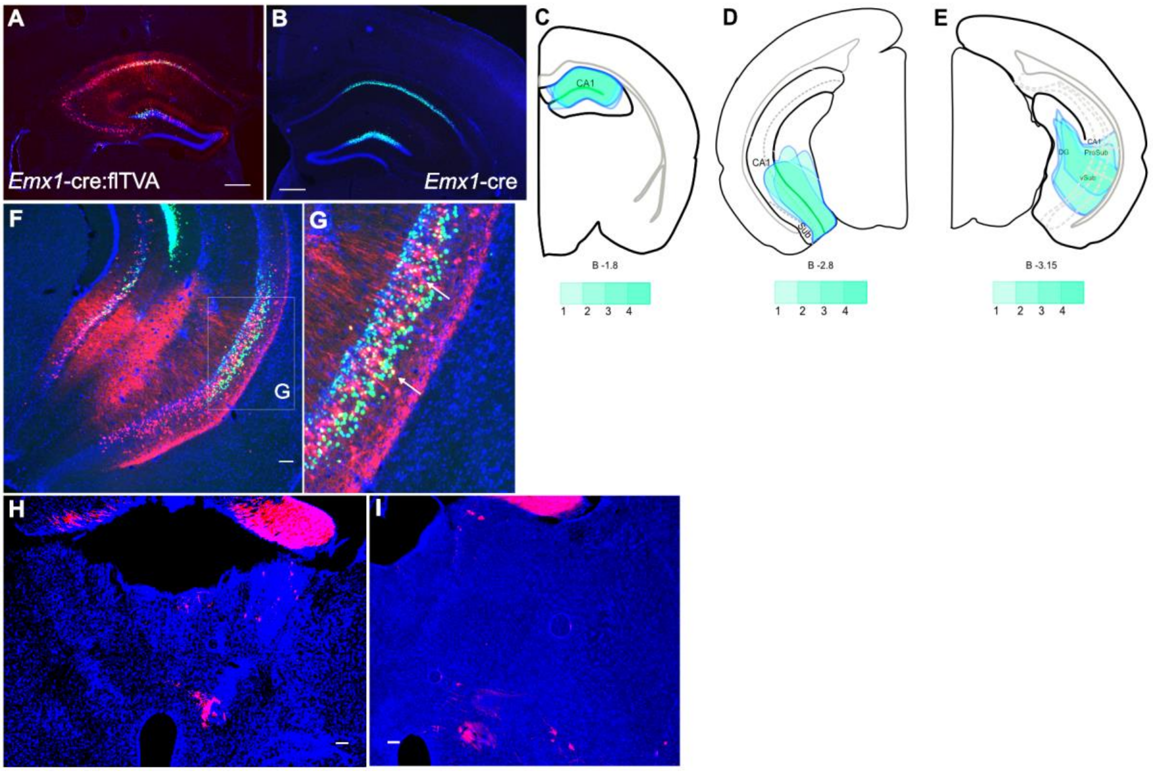


**Figure S5: control injections for retrograde monosynaptic tracing. A**, representative example of injection site for retrograde labelling viruses in *Emx1-cre:*TVA mouse. Both green (transduced by the helper AAV virus) and red (transduced by the pseudotyped rabies virus) cells can be seen. **B**, in a control mouse (*Emx1-cre*) without TVA, the pseudotyped rabies virus was unable to enter CA1 pyramidal cells (n=6 mice). Scale bar represents 100 microns. **C – E**, schematic images showing spread of viral representative spreads of virus for injections in the dCA1 (**C**), vCA1 (**D**) and vSub (**E**) regions. Each figure is a composite of 4 injections, with the darker colour indicating more mice. **F & G,** representative example of injection on low (**F**) and high (**G**) magnification of injections that spread between vCA1, prosubiculum and vSub. **H & I**, retrogradely-labelled neurons were present in NRe only in mice with starter cells located in prosubiculum and subiculum, confirming our observations from anterograde optogenetics experiments (figure 1).

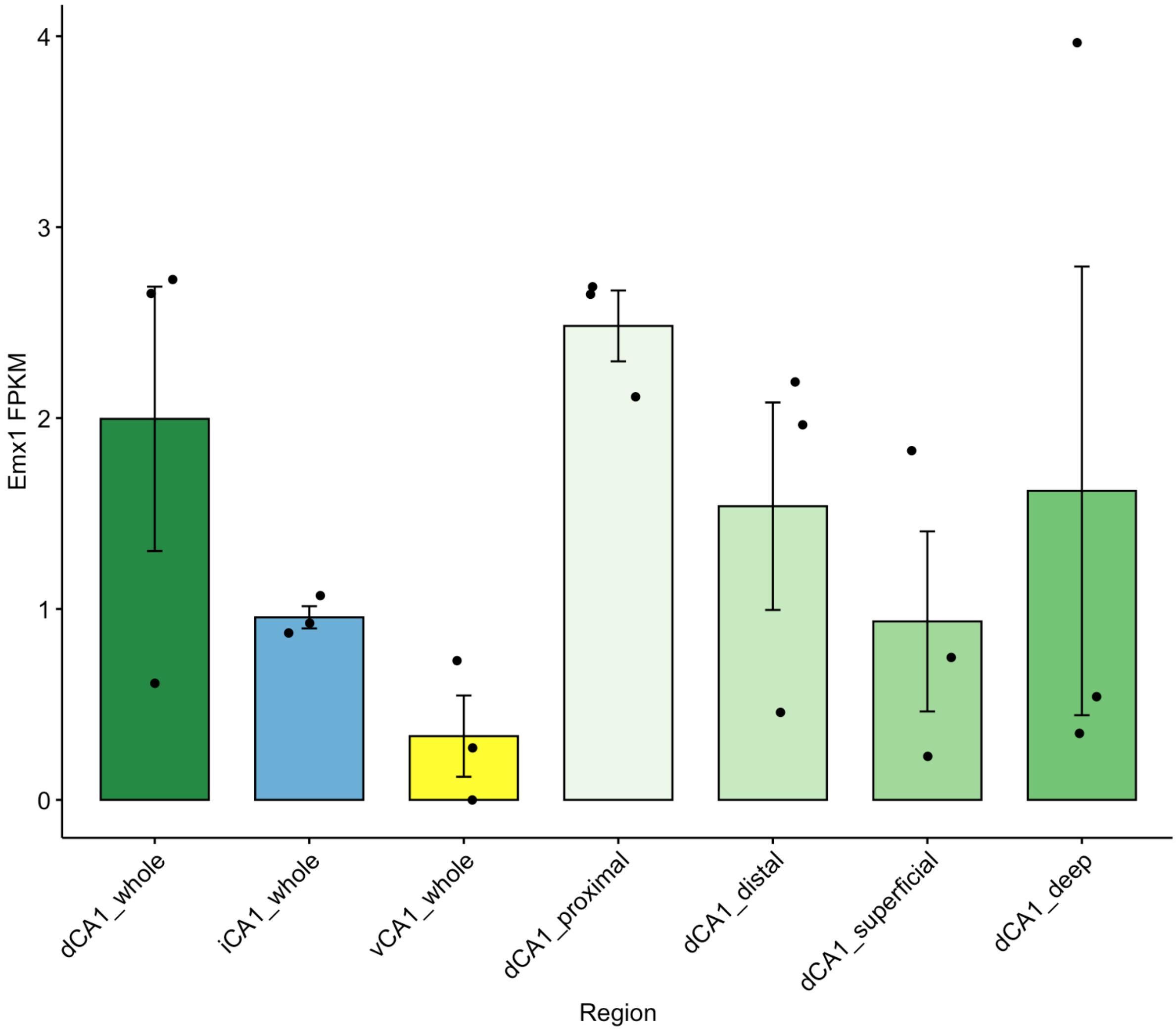


**Figure S6: *Emx1* is expressed throughout the hippocampus.** Secondary analysis of Cembrowksi *et al., 2016, doi: 10.1016/j.neuron.2015.12.013)* revealed that *Emx1* expression is ubiquitous throughout hippocampus, albeit at (non-significantly) lower levels in ventral CA1 compared with dorsal or intermediate CA1.
